# Natural killer cell-mimic nanoparticles can actively target and kill acute myeloid leukemia cells

**DOI:** 10.1101/2022.09.09.507236

**Authors:** Hojjat Alizadeh Zeinabad, Wen Jie Yeoh, Mihai Lomora, Yara Banz, Carsten Riether, Philippe Krebs, Eva Szegezdi

## Abstract

Natural killer (NK) cells are effector lymphocytes of the innate immune system which play a crucial role in recognizing and killing emerging tumor cells. However, as the tumor evolves, it develops mechanisms to inactivate NK cells or hide from them. Here, we engineered a modular nanoplatform that acts as NK cells (NK cell-mimics), carrying the tumor-recognition and death ligand-mediated tumor-killing properties of an NK cell, yet without being subject to tumor-mediated inactivation. In particular, NK cell mimic nanoparticles (NK.NPs) incorporate two key features of activated NK cells: cytotoxic activity via the death ligand, tumor necrosis factor-related apoptosis-inducing ligand (TRAIL), and an adjustable tumor cell recognition feature based on functionalization with the NK cell Fc-binding receptor (CD16, FCGR3A) peptide, enabling the NK.NPs to bind antibodies targeting tumor antigens. NK.NPs showed potent *in vitro* cytotoxicity against a broad panel of cancer cell lines. Upon functionalizing the NK.NPs with daratumumab, a clinically used antibody specific for the CD38 protein expressed by AML cells, NK.NPs effectively targeted and eliminated patient-derived acute myeloid leukemia (AML) blasts and leukemia-initiating cells as well as CD38-positive AML cells *in vivo*, in a disseminated AML xenograft system. Specifically, NK.NPs lead to a significant reduction of AML burden in the bone marrow, spleen, and peripheral blood compared to non-targeted TRAIL-functionalized liposomes. Taken together, these findings demonstrate that NK.NPs are effective in mimicking NK cells’ antitumorigenic function and thereby underline their use as therapeutic tools.

## Introduction

Natural killer (NK) cells play a crucial role in recognizing and killing emerging tumor cells. Unlike T cells, as innate immune cells, NK cells can attack unwanted cells, including tumor cells without prior antigen sensitization and priming by antigen-presenting cells. This is because instead of an immunoglobulin-like antigen receptor, NK cells use an array of activating receptors that recognize a range of cell stress markers. Of these receptors, CD16 stands out for its therapeutic potential. CD16 (Fc gamma receptor III, FCGR3A or FcgRIII) binds to the fragment crystallizable constant region (Fc region) of antibodies and can thus recognize antibody-targeted cells and particles (opsonization), a mechanism that can be utilized for tumor cell recognition.^1–3^

Similar to other activating NK cell receptors, antibody binding by CD16 initiates target cell killing which takes place via two major pathways, i.e. release of cytolytic granules containing pore-forming proteins, such as perforin and proteolytic enzymes like granzyme B, or through binding of death receptors to cognate death ligands on target cells, namely Fas ligand and TNF-related apoptosis-inducing ligand (TRAIL).^2^ However, some tumor cells can escape immune recognition and develop mechanisms to hide from or to inactivate NK cells. For instance, tumor cells can shed NKG2D ligands, which mask this activating NK receptor. Cancer cells can also escape from NK cell-mediated killing by producing immune-suppressive factors, such as prostaglandin E2, interleukin-10 and transforming growth factor-β (TGFβ).^4^

Death ligands expressed by effector immune cells have been considered as anti-cancer therapeutics since their discovery. Of these death ligands, TRAIL has the highest potential as an anti-cancer therapeutics due to its low off-target toxicity.^5^ Preclinical studies have demonstrated the potent anti-tumor activity of soluble, recombinant human TRAIL (sTRAIL) protein with minimal to no side effects. Yet, the results of the subsequent clinical trials showed lack of efficacy, which was attributed to limited cytotoxicity of TRAIL when used as a soluble, recombinant protein, short circulatory half-life time and insufficient tumor accumulation. Nevertheless, TRAIL shows strong tumoricidal activity when expressed by effector immune cells.^6,7^ Thus, generating a TRAIL-based therapeutics that mimics effector immune cells can address the limitations of sTRAIL and could thus lead to a well-tolerated and effective cancer treatment.^8^ Nanoparticles (NPs) offer such a solution based on previous studies which demonstrated that conjugation of TRAIL to nanoparticles can overcome some limitations of sTRAIL, such as increasing its cytotoxic activity via stabilizing its oligomeric structure and increasing its serum half-life time.^9^

Most importantly, it has been shown that one of the major challenges in developing a system for active targeting of tumor cells is the heterogeneity of tumors across patients (interpatient heterogeneity), even within the same tumor type.^10^ Thus an effective targeting system requires adjustability to enable tailoring the treatment to specific patient cohorts.

Here, we have developed a novel, NK cell-mimic NP (NK.NP) that carries essential NK cell functions and kills cancer cells via TRAIL. The designed lipid-based NK.NP encompasses CD16-dependent cell recognition and TRAIL-mediated killing function of NK cells, but unlike NK cells and effector immune cells in general, it cannot be inactivated via immune editing by tumors. The NK.NPs could efficiently target and kill acute myeloid leukemia (AML) *in vitro, ex vivo* as well as *in vivo* without detectable systemic toxicity. These results show that NK.NPs have an excellent potential to be developed into off-the-shelf, easy to generate and potent immunotherapeutic agents against a broad range of cancers.

## Methodology

### Materials and reagents

All lipids were purchased from Avanti. All chemicals and reagents used were purchased from Sigma-Aldrich unless otherwise stated.

### Liposome formation and labelling

Monodisperse liposomes comprised of 1,2-distearoyl-sn-glycero-3-phosphocholine (DSPC), 1,2-distearoyl-sn-glycero-3-phosphoethanolamine-N-[methoxy(polyethylene glycol)-2000] (DSPE-PEG), 1,2-distearoyl-sn-glycero-3-phosphoethanolamine-N-[PDP(3-(2-pyridyldithio) propionate) polyethylene glycol-2000] (DSPE-PEG-PDP) were prepared using thin-film hydration method followed by extrusion. Briefly, DSPC, DSPE-PEG and DSPE-PEG-PDP dissolved in chloroform were mixed at a molar ratio of 9:0.5:0.5 and the chloroform was evaporated under a stream of nitrogen to form a film in a round bottom flask and kept under vacuum overnight. Next day, the dry film was rehydrated with Tris buffer (50 mM, pH 7.4) in a rotary evaporator at 65°C for one hour to self-assemble phospholipids into liposomes (5 mg lipid/ml). For *in vivo* studies, liposomes were prepared in potassium phosphate buffer (PBS, BioSciences). The polydisperse multilamellar liposomes were extruded through a 200 nm pore-size polycarbonate filter 11 times and through a 100 nm polycarbonate membrane 17 times using an Avanti Mini Extruder (Avanti Polar Lipids, AL, USA) to obtain small unilamellar liposomes. Non-PEGylated liposomes were synthesized with the same method using 1,2-distearoyl-sn-glycero-3-phosphocholine (DSPC) and 1,2-distearoyl-sn-glycero-3-phosphoethanolamine (DSPE) in a molar ratio of 9:1. For CM-DiI fluorescent membrane tracker-labelling, liposomes were incubated with 150 nM 3H-Indolium, 5-[[[4-(chloromethyl)benzoyl]amino]methyl]-2-[3-(1,3-dihydro-3,3-dimethyl-1-octadecyl-2H-indol-2-ylidene)-1-propenyl]-3,3-dimethyl-1-octadecyl-, chloride (CM-DiI, Invitrogen) for 20 minutes at 37°C followed by centrifugation at 30,000 x g for one hour at 4°C to remove excess CM-DiI.

### Expression and purification of Strep-tagged TRAIL

The extracellular fragment of TRAIL (fragment: amino acids 114-281) was cloned into a pET-51b (+) plasmid with an N-terminal Strep-tag II tag. The expression vector was transformed into BL21 *E. coli* and cultured in Luria Broth (LB) medium containing 1% (w/v) glucose and 100 μg/ml ampicillin in a shaking incubator at 37 °C until reached an optical density (OD600) of ∼0.7. 1 mM isopropyl β-D-1-thiogalactopyranoside (IPTG, Roche) and 100 μM ZnSO_4_ were added to the culture and bacterial culture was continued at 28°C for 5 hours (h) to induce Strep-TRAIL expression as previously described.^11^ Subsequently, bacteria were harvested and resuspended in lysis buffer (50 mM Tris, 150 mM NaCl, 10% glycerol, 7 mM 2-mercaptoethanol, pH 8.0) and lysed with a probe sonicator (25% amplitude, 12 cycles, 20 seconds on, 45 seconds off on ice) followed by centrifugation. The supernatant was subjected to purification using a StrepTrap™ HP affinity column (IBA Lifesciences) on an AKTA FPLC system and eluted with elution buffer (50 mM Tris, 150 mM NaCl, 10% glycerol, 3.5 mM DTT, 20 μM ZnSO_4_, 2.5 mM desthiobiotin, pH 7.4). The purified protein was quantified by bicinchoninic acid assay (BCA) (Thermo Fisher Scientific) and nanodrop (Thermo Fisher Scientific) and stored at −80°C until use.

### Analytical SDS-PAGE and Western Blotting

Samples collected from stages of Strep-TRAIL expression and purification were analyzed by sodium dodecyl sulfate-polyacrylamide gel electrophoresis (15%) and Western blotting. Yield and purity were assessed by staining the gel with Coomassie blue, and expression of Strep-TRAIL determined after transferring the proteins from the gel onto nitrocellulose membrane (iBlot, Invitrogen). After blocking in 5% milk in PBS/0.05% Tween-20, the membrane was incubated with a mouse monoclonal Strep-tag antibody (1:2000 dilution, Qiagen), followed by incubation with a secondary antibody conjugated to horse-radish peroxidase (HRP, 1:3,000 dilution, Pierce). Enhanced chemiluminescence (ECL) reagent (Thermo Fisher Scientific) was applied to visualize protein bands on an X-ray film (Agfa).

### Preparing NK cell-mimic nanoparticles

A 3-(2-pyridyldithio) propionate (PDP) group-conjugated DSPE-PEG (DSPE-PEG-PDP) was used to covalently link streptavidin to the surface of the liposome, enabling liposome functionalization with Strep-tagged and biotinylated molecules in an oriented manner. The PDP group was reduced by incubating the liposomes with 20 mM dithiothreitol (DTT) for 30 minutes at room temperature (RT). After PDP-reduction, DTT was removed by ultracentrifugation at 30,000 x g for one hour at 4°C and resuspension in Tris buffer (50 mM, pH 7.4). The thiolated liposomes were then incubated with maleimide-activated streptavidin (STV, ThermoFisher Scientific) at a molar ratio of 1:10 (STV:PDP-PEG-DSPE) with gentle stirring at 4°C for 16 h and then the mixture was centrifuged again to remove unbound STV. To verify liposome surface modification with streptavidin, binding of Atto 488-biotin (ATTO-TEC) was measured with flow cytometry. Streptavidin conjugated liposomes (LP/STV) were then resuspended in Tris buffer (50 mM, pH 7.4) and functionalized with 50 μg/ml Strep-TRAIL or 30 μg/ml Fc gamma receptor peptide/immunoglobulin-binding peptide (FCP, peptide sequence: FNMQQQRRFYEALHDPNLNEEQRNAKIKSIRDD GGG-Lys-Biotin, ProteoGenix, France)^12^ for one hour at RT to obtain liposomes functionalized with TRAIL (LP/STV/TRAIL) or FCP (LP/STV/FCP), respectively. Liposomes were ultracentrifuged again to remove unbound TRAIL and FCP, and the pellets were resuspended in Tris buffer. To confirm functionalization with Strep-TRAIL and FCP, binding of APC-conjugated anti-TRAIL antibody (APC-TRAIL-Ab, BD Biosciences) and FITC-tagged anti-CD123 antibody (FITC-Ab) was used. The fluorescence intensity proportional to antibody binding was measured with flow cytometry.

The surface of LP/STV/FCP was further modified with monoclonal anti-CD38 antibody (daratumumab, Darzalex, Janssen) binding to the FCP by incubating the liposomes with 200 μg/ml antibody for one hour at RT (LP/STV/FCP/Ab). For the removal of excess antibodies, liposomes were ultracentrifuged. Finally, to obtain NK.NPs, TRAIL- and FCP+antibody-functionalized liposomes were combined by incubating their equimolar mix for 2 h at RT with brief vortexing every 15 minutes, followed by extrusion through a 200 nm polycarbonate membrane 11 times at 37°C. To concentrate the NK.NPs for *in vivo* study, after final extrusion, they were ultracentrifuged and resuspended in PBS to obtain 100 μg /ml TRAIL NK.NPs. To determine the binding affinity of FCP to antibodies, a serial dilution of the FCP was conjugated to the bottom of wells of a streptavidin-coated 96-well plate (Thermo Fisher Scientific) for 60 minutes at RT. After removing excess FCP with three washes with wash buffer (25 mM Tris, 150 mM NaCl; pH 7.2), the plate was incubated with a FITC-Ab for 60 minutes at RT. Excess, unbound FITC-Ab was removed in three washes with wash buffer and the fluorescence intensity of the FCP-bound FITC-Ab determined using a VICTOR plate reader (Perkin Elmer, US). To examine the targeting ability of liposomes functionalized with antibody, LP/STV/FCPs were incubated with anti-CD123 antibody and labelled with CM-DiI (DiI/LP/STV/FCP/CD123 Ab). CD123-positive KG1a cells were labelled with 1 μM final concentration of CellTracker™ deep red dye (Thermo Fisher Scientific) as per the manufacturer’s instructions to allow separation of KG1a cells from liposomes during analysis and the cells were incubated with DiI/LP/STV/FCP/CD123 Ab for 90 minutes, followed by measuring fluorescence intensity by flow cytometry.

### Cell culture

HeLa, SW1990 and MDA-MB-231 cells were grown in DMEM medium, and primary human fibroblasts were cultured in low glucose DMEM medium (with 1000 mg/l glucose, L-glutamine, and sodium bicarbonate). K562, MM1.S, NCI-H929, Colo 205, U2OS, BXPC-3, ML-1, ML-2, Kasumi-1, and KG1a cells were grown in RPMI-1640 medium and OCI-AML2 cells were grown in alpha-MEM medium. All media were supplemented with streptomycin (100 μg /ml), penicillin (100 U/ml) and 10% Hyclone fetal bovine serum (FBS, Thermo Fisher Scientific). The medium for HeLa cells was further supplemented with 1% MEM amino acids solution, and L-glutamine (2 mM). Also, the medium for OCI-AML2, ML-1, ML-2, Kasumi-1, KG1a, and Colo 205 cells was further supplemented with L-glutamine (2 mM), sodium pyruvate (1 mM), and 1% RPMI-1640 amino acids solution and the cells were grown in a humidified incubator (5% CO_2_, 37°C). Primary AML samples were obtained from Blood Cancer Biobank Ireland (Galway, Ireland). Peripheral blood samples from healthy donors were obtained from consented healthy volunteers. Mononuclear cells (MNC) were separated from peripheral blood, and bone marrow aspirates samples using Ficoll gradient separation as described before.^13^ Cells were cultured in RPMI-1640 medium supplemented with streptomycin (100 μg /ml), penicillin (100 U/ml), 10% FBS, 2 mM L-glutamine, and 1 mM sodium pyruvate in a humidified incubator (5% CO_2_, 37°C). KHYG-1 cells were grown in RPMI-1640 medium (with L-glutamine and sodium bicarbonate) supplemented with 10% heat-inactivated fetal bovine serum, streptomycin (100 μg /ml), penicillin (100 U/ml), and interlukin-2 (100 U/ml) (PeproTechTM). The monocytic, human acute monocytic leukemia cell line, THP-1, was cultured in RPMI-1640 supplemented with 10% FBS, streptomycin (100 μg /ml), penicillin (100 U/ml) under a humidified 5% (v/v) CO_2_ atmosphere at 37°C.

### Colony forming assay

Primary AML cells were isolated from patient-derived xenografts (PDX) in NSG mice, and cells were seeded in round bottom 96-well plates. The cells were then treated with 10 μl of LP/STV as a negative control, 250 ng soluble recombinant human TRAIL (sTRAIL), 25-250 ng of TRAIL conjugated to liposomes (LP/STV/TRAIL) or CD38-NK.NPs overnight. After treatment, the cells were transferred into methylcellulose enriched media (R&D Systems) and cultured for 11-13 days after which the formed stem cell-derived colonies were enumerated under a light microscope.

### Cell viability assay

Induction of cell death in cancer cell lines was measured with Annexin V assay, and cell death of primary cells (AML patient samples and healthy donor samples) was measured with the viability dye, TO-PRO 3 (Invitrogen). Suspension cells (PBMC, K562, MM1.S, NCI-H929, Colo 205, U2OS, KG1a, OCI-AML2, ML-1, ML-2 and Kasumi-1 cells) were harvested by pipetting while adherent cells (HeLa, SW1990, BXPC-3, MDA-MB-231 and healthy primary human fibroblasts) were harvested by trypsinization. Cells were collected by a pulse spin at 14,000 rpm for 10 seconds, the supernatant removed, and the cells were resuspended in 50 μl Annexin V buffer (10 mM HEPES/NaOH, pH 7.5, 2.5 mM CaCl2,140 mM NaCl) containing 1.5 μl Annexin V-FITC or -APC (Immunotools) for 15 minutes on ice in the dark. For TO-PRO 3 staining, cells were incubated with 50 μl PBS containing 1 μl TO-PRO 3 (50 μM) for 15 minutes and volume was increased to 200 μl with PBS buffer before analysis with flow cytometry. The percentages of Annexin V-positive cells and TO-PRO 3 positive cells were determined with flow cytometry.

### NK cell-mediated cellular cytotoxicity assay

KHYG-1 NK cells were labelled with the green cell tracker CFSE (carboxyfluorescein succinimidyl ester, 2.5 μM, Biolegend) as per the manufacturer’s instructions to allow separation of NK cells from AML cells during analysis. CFSE-labelled KHYG-1 cells (effector cell) and AML cells (target cells) were co-cultured at target to effector (T:E) ratios of 1:1, 1:2.5 and 1:5 for 24 h after which the cells were collected and the percentage of AML cells killed was determined with Annexin V-APC labelling as described above. For analysis, the effector cells were excluded based on CFSE positivity and the percentage of Annexin V-APC positive dead cells was determined within the AML population.

### THP-1 macrophage differentiation and liposome phagocytosis assay

THP-1 monocytes (80,000 cells/well) were differentiated into macrophages by treatment with 100 ng/ml phorbol 12-myristate 13-acetate (PMA). After 24 h PMA treatment, supernatants were replaced with fresh media and the generated macrophages were incubated with a dosage of PEGylated, and non-PEGylated liposomes loaded with CM-DiI for 3 h. Subsequently, supernatants, containing non-phagocytosed liposomes, and the cells with engulfed liposomes (after detachment with trypsin/EDTA) were collected separately. The cells were labelled with CD45-APC and re-pooled with the supernatants before FACS analysis to quantify phagocytosed liposomes.

### Immunophenotyping

Cells were harvested by centrifugation at 14,000 rpm for 10 seconds, washed with 1% BSA in PBS, blocked in the same buffer and incubated with fluorescently labelled antibodies against DR4, DR5, CD38 and CD45 for 30 minutes on ice with intermittent mixing. Excess antibodies were removed by washing with PBS/1%BSA and antibody-fluorescence intensity was measured with flow cytometry (FACS SORP LSR II, FACS Canto A or FACS Canto II flow cytometers, BD Bioscience, San Diego, USA). Data were analyzed using Flowjo software or FCS Express V3 software.

### Dynamic light scattering

The liposomes’ hydrodynamic diameter and zeta potential were determined using Zetasizer Nano ZS (Malvern) or Litesizer 500 (Anton Paar) at room temperature. Liposomes were diluted to approximately 200 μg/ml in Tris buffer (50 mM, pH 7.4) to a final volume of 1 ml, and then size distribution was measured. Mean liposome size in three forms, intensity (Z-average), number and volume averages, were determined using the Zetasizer software. For measuring surface charge, liposomes were diluted to 200 μg/ml and the surface charge measurements were conducted as average zeta potential of the liposomes at 25°C.

### Transmission electron microscopy

The morphology and size of the liposomes were characterized with a Hitachi Transmission Electron Microscope (Hitachi H-7500). Liposomes were diluted to approximately 2 mg/ml, and then 3 μl was deposited onto 200-mesh copper grids coated with a Formvar support film and dried in a chemical hood for 5 minutes. The grids were analyzed and photographed at a magnification of 30,000x and an acceleration voltage of 75 kV.

### *In vivo* studies in disseminated AML xenograft mouse model

5-6 weeks old female immune deficient NOD SCID gamma (NSG) mice were purchased from Charles River Laboratories (Germany) and subsequently bred in-house in pathogen-free facilities. The experimental protocols conformed to the ethical guidelines of the Swiss Federal regulations and were approved by the Cantonal Veterinary Office of Bern, Switzerland (license, BE-13-2021). During all experiments, mice were observed and weighed daily, and their behavior was monitored over time. At the end of each experiment, mice were euthanized using CO_2_ inhalation. Spleens and femurs were extracted. Spleens were smashed using the plunger of a 5 ml syringe and were filtered through a 70 μm cell strainer (VWR, China). Femurs were cleaned, the heads were cut with surgical scissors and the bone marrow was flushed out with PBS and passed through a 40 μm cell strainer (FALCON, USA) to remove debris. To isolate cells from peripheral blood, spleen, and femurs, collected samples were spun at 500 x g for 5 minutes. Pellets were resuspended in 1 ml ACK (Ammonium-Chloride-Potassium) lysis buffer (150 mM ammonium chloride, 10 mM potassium bicarbonate, and 0.1 mM EDTA in ultrapure water) and were incubated for 5 minutes at RT to lyse the red blood cells. Next, samples were washed with cold PBS and incubated with a mixture of anti-mouse CD45 antibody (mCD45, Invitrogen), anti-human CD45 antibody (hCD45, Bioligand) and a live/dead fixable blue dead cell stain (Invitrogen) for 30 minutes at 4ºC. Then samples were washed two times with PBS, followed by measuring the fluorescence intensity by flow cytometry.

To establish a disseminated AML xenograft, 8-10 weeks old female NSG mice were administered with 1×10^6^ OCI-AML2 cells (suspended in 200 μl PBS) via the tail vein. Tumor burden was monitored three-daily by collecting peripheral blood from the tail vein and measuring the frequency of human CD45^+^ cells. At the end of experiments, mice were euthanized using CO_2_ inhalation and disease burden assessed as described above.

To assess the toxicity of NK.NPs, 8–9-week-old healthy female NSG mice were administered with 500 μl of either NK.NPs or PBS (as control) intra-peritoneally (i.p.) twice, with a 48 h gap. Signs of toxicity during the treatments were monitored. Mice were euthanized 48 hours after the second injection (at 96 h), and their liver, spleen and kidneys were extracted and weighed. Percentage of body weight changes were calculated. Peripheral blood was collected and subjected to complete blood cell count (CBC) and plasma biochemical analysis. Plasma samples were obtained by incubating the collected blood at RT for 30 minutes, followed by centrifugation in SST tubes (BD Microtainer) at 15,000 xg for 90 seconds and the levels of aspartate aminotransferase (AST), alanine aminotransferase (ALT), glucose, albumin, blood urea nitrogen (BUN) and creatinine were determined.

In order to investigate the efficacy of NK.NPs, first, AML cells were engrafted into 8-10 weeks old female NSG mice as described above. Then, mice were randomized into four groups, and 500 μl of either sTRAIL, naked liposomes, LP/TRAIL or NK.NPs were injected (i.p) from day 4 of AML post implantation by i.p injection in a total of five dosages every other day. Body weight and behavior were monitored every day. Tumor burden was monitored three-daily as described above. Mice were euthanized 14 days after AML cell injection, and their liver, spleen, tibia, and femurs were extracted. Spleens and livers were weighed, and spleen size was measured. Femurs and spleens were subjected to cell isolation for immunophenotyping and tibias were fixed in formalin for 24 h for histopathological evaluation followed by staining with hematoxylin and eosin (H&E). Frequencies of AML blasts in tissue sections were quantified by a board-certified hemato-pathologist (Y.B.).

## Results

### Design, synthesis and characterization of nanoparticles

NK.NPs were designed to possess the key NK cell effector functions of recognizing and killing malignant cells. The base-formulation of the NP was designed to meet the standard criteria for nanoparticles, which are to be negatively charged and between 10 – 150 nm in size to avoid filtration by the kidneys, but able to pass through the endothelial barrier.^14,15^ Liposome formulation was chosen due to its biocompatibility, cell-like characteristics and ability to penetrate through vessel walls.^16^

To meet these requirements, liposomes composed of 1,2-distearoyl-sn-glycero-3-phosphocholine (DSPC), 1,2-distearoyl-sn-glycero-3-phosphoethanolamine-N-[(polyethylene glycol)-2000] (DSPE-PEG2000), 1,2-distearoyl-sn-glycero-3-phosphoethanolamine-N-[PDP(3-(2-pyridyldithio) propionate) polyethylene glycol-2000] (DSPE-PEG-PDP) were generated. Dynamic light scattering (DLS) and transmission electron microscopy (TEM) studies confirmed that the generated liposomes are monodispersed with a polydispersity index (PDI) of 0.133 and an average hydrodynamic size of 98.12 nm (Figure 1).

**Figure 1.**
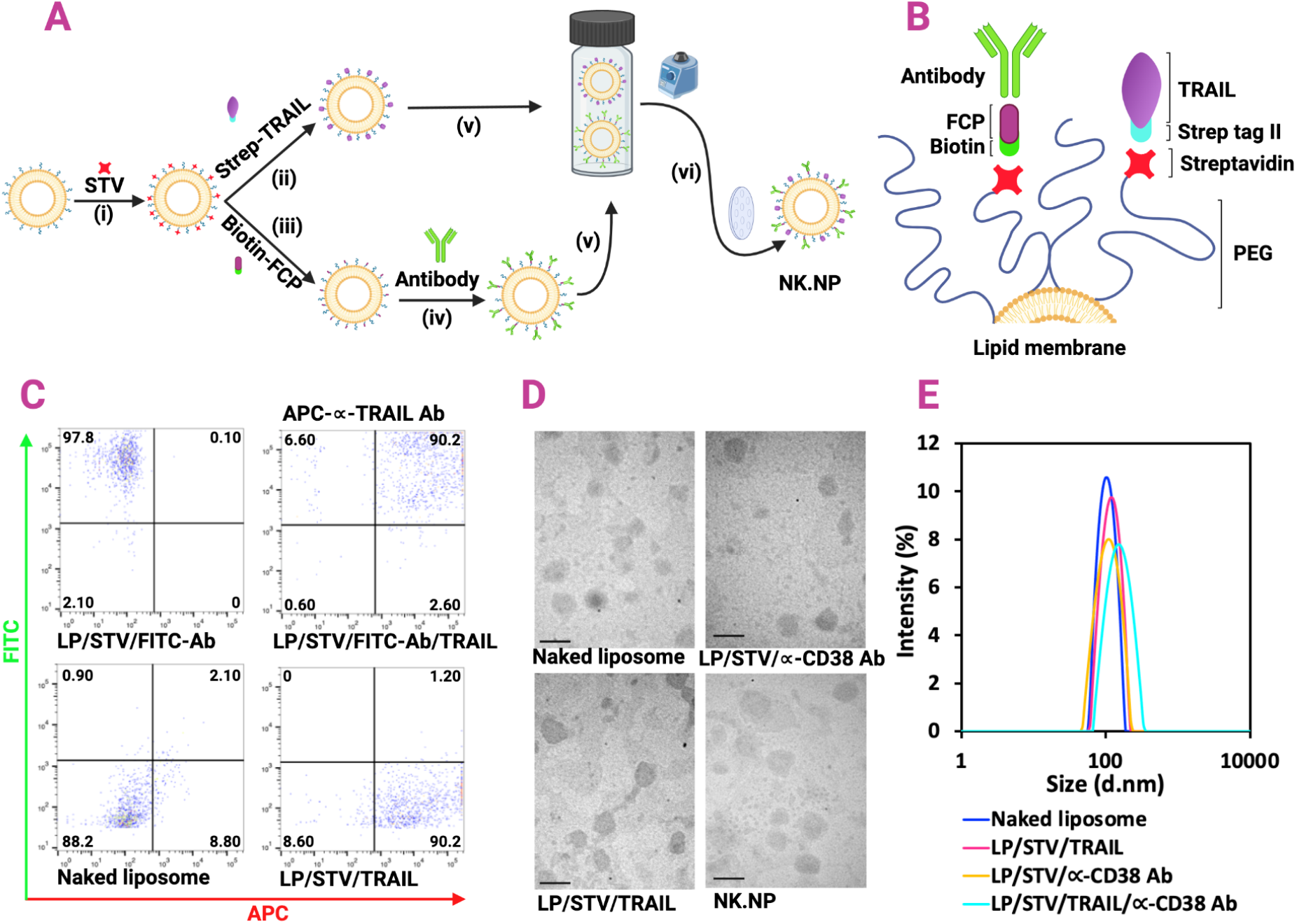
Generation of natural killer cell-mimic nanoparticles. (A) Schematic illustration of the NK cell-mimic nanoparticle (NK.NP) fabrication process. (i) Liposome surface modification with streptavidin (STV) (ii) followed by functionalization with Strep-TRAIL or (iii) biotin-conjugated immunoglobulin binding peptide (FCP) and (iv) therapeutic monoclonal antibody. (v) Fusion of TRAIL-functionalized liposomes (LP/STV/TRAIL) with antibody-functionalized liposomes (LP/STV/αCD38 Ab) at room temperature with interval vortex (vi) followed by passing through polycarbonate membrane using an extruder to generate NK.NPs. (B) Schematic of the composition of NK.NPs. (C) Representative flow cytometry plots showing the successful functionalization with FCP, Strep-TRAIL or both detected with FITC-conjugated antibody (LP/STV/FITC-Ab for FCP) and APC-tagged anti-TRAIL antibody (APC-TRAIL Ab for Strep-TRAIL). (D) Representative electron micrographs and (E) DLS measurements of liposome structures (scale bar = 200 nm).

Soluble TRAIL protein has an approximately 100-1,000-fold lower cytotoxic activity that membrane-bound TRAIL and in resistant tumor cells, it may even have a pro-tumorigenic activity by promoting migration and thus potentially driving cancer metastasis^17^. To overcome this, we presented TRAIL in its native, membrane-bound orientation and conformation on the NK.NPs, thus replicating the potent activity of surface-expressed TRAIL of NK cells. To achieve this, sTRAIL was tagged with Strep-tag II on its cytosol-facing N-terminus (Strep-TRAIL), enabling its conjugation in the correct orientation to streptavidin (STV) moieties on the liposome surface. Since STV has 4 Strep-tag/biotin binding sites, TRAIL conjugation to STV also resulted in TRAIL clustering, while the lipid bilayer of the liposome enabled lateral movement of the conjugated TRAIL facilitating high-order aggregation upon receptor binding (capping) required for apoptosis induction.^18^

Strep-TRAIL was expressed in BL21 *Escherichia coli* and purified using affinity chromatography. Coomassie blue gel staining and western blotting confirmed high purity (Supplementary Figure 1A-C) and FPLC chromatogram confirmed the predominant single TRAIL species which did not form aggregates or contained impurities (Supplementary Figure 1C). The biological activity of Strep-TRAIL was confirmed by testing its cytotoxic potential on the TRAIL-sensitive colorectal cancer cell line, Colo205 that showed a comparable cytotoxicity of Strep-TRAIL to untagged sTRAIL (Supplementary Figure 1D, 1E).

Next, the surface of the liposomes was modified with maleimide-conjugated STV via a thiol-linkage to create an attachment site for Strep-TRAIL. Functionalization of the liposomes with STV was confirmed by measuring the ability of the liposomes to bind Atto 488-biotin (Supplementary Figure 2A, 2B). The STV-functionalized liposomes (LP/STV) were then conjugated with Strep-TRAIL, which was confirmed by detecting TRAIL on the LP/STV surface with anti-TRAIL antibody (APC-TRAIL Ab, Supplementary Figure 3B) and by measuring the biological activity of LP/STV-bound TRAIL (LP/STV/TRAIL) against Colo205 cells (Supplementary Figure 3C).

To ensure saturation of the TRAIL-binding sites on the liposomes, LP/STV NPs were functionalized with increasing concentrations of Strep-TRAIL. The cytotoxic potential of the generated LP/STV/TRAIL as well as the excess, unbound Strep-TRAIL in the reaction supernatant, were then measured against Colo205 cells. Conjugation with 30 μg/ml Strep-TRAIL achieved maximum killing of Colo205 cells. Simultaneously, the supernatant from the conjugation reactions with more than 50 μg/ml Strep-TRAIL started to show cytotoxicity, indicating saturation of liposomes with TRAIL (Supplementary Figure 4). Thus, 50 μg/ml Strep-TRAIL was chosen as the optimal concentration to functionalize the liposomes for the rest of the experiments.

To assess the biological activity of TRAIL-conjugated liposomes (LP/STV/TRAIL), their ability to escape phagocytosis and potency to trigger cell death in comparison to sTRAIL was assessed. LP/STV/TRAIL was generated with incorporation of polyethylene glycol (PEG) on the lipids to reduce phagocytosis. As expected, the presence of PEG significantly reduced the engulfment of liposomes by THP-1-derived macrophages (Supplementary Figure 5), suggesting a longer bioavailability *in vivo*.

The tumoricidal potency of LP/STV/TRAIL was then tested on a broad panel of cancer cell lines with ranging TRAIL sensitivities, including leukemia (K562 and KG1a), myeloma (sMM1.S and NCI-H929), colorectal-(Colo205), breast-(MDA-MB-231), cervical-(HeLa), pancreatic-(BXPC-3 and SW1990) and osteosarcoma-(U2OS) cancer cells. The cells were treated with either sTRAIL or LP/STV/TRAIL for 24 h, and cell death was quantified with Annexin V staining. LP/STV/TRAIL showed a superior cytotoxic activity over sTRAIL in in the large majority of investigated cancer cell lines (Figure 2A-J) with the exception of KG1a cells (Figure 2K). Importantly, LP/STV/TRAIL was not toxic to non-malignant cells, including primary human fibroblasts and peripheral blood leukocytes (Figure 3).

**Figure 2.**
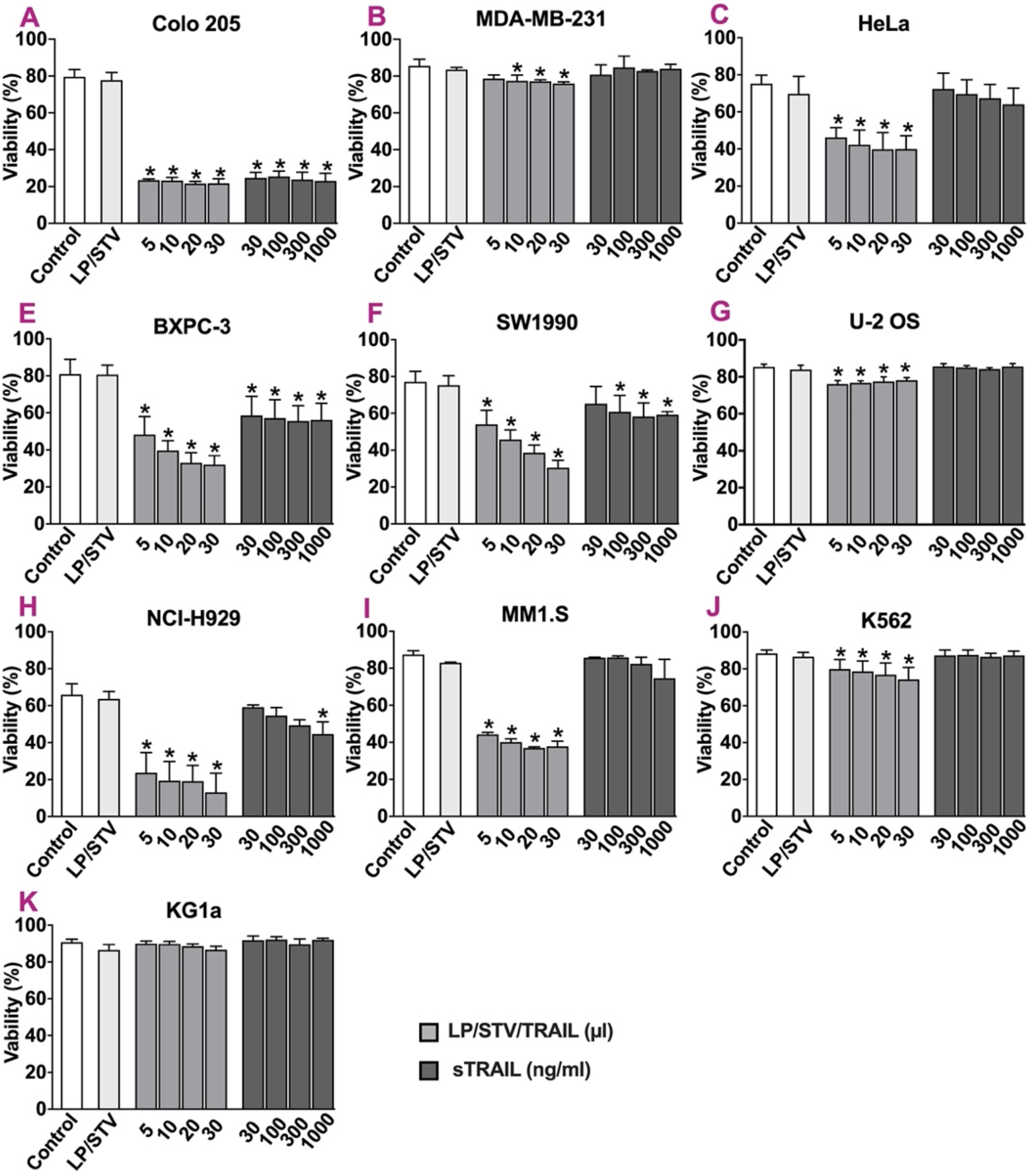
TRAIL-conjugated liposomes have higher tumoricidal potential compared to soluble TRAIL. Cancer cell Colo205 (A), MDA-MB-231 (B), HeLa (C), BXPC-3 €, SW1990 (F), U-2 OS (G), NHI-H929 (H), MM1.S (I), K562 (J), and KG1a (K) were incubated with 5-30 μl of TRAIL-conjugated liposomes (LP/STV/TRAIL), 30 μl streptavidin-conjugated liposomes (LP/STV) as negative control and 30-1000 ng/ml soluble recombinant human TRAIL (sTRAIL) for 24 h. Apoptosis was quantified using Annexin V-FITC staining. Bars represent the mean percentage of live cells ± standard deviation of at least three independent repeats. The statistical significance was evaluated using one-way ANOVA followed by Tukey’s test with multiple comparison. *P<0.05.

**Figure 3.**
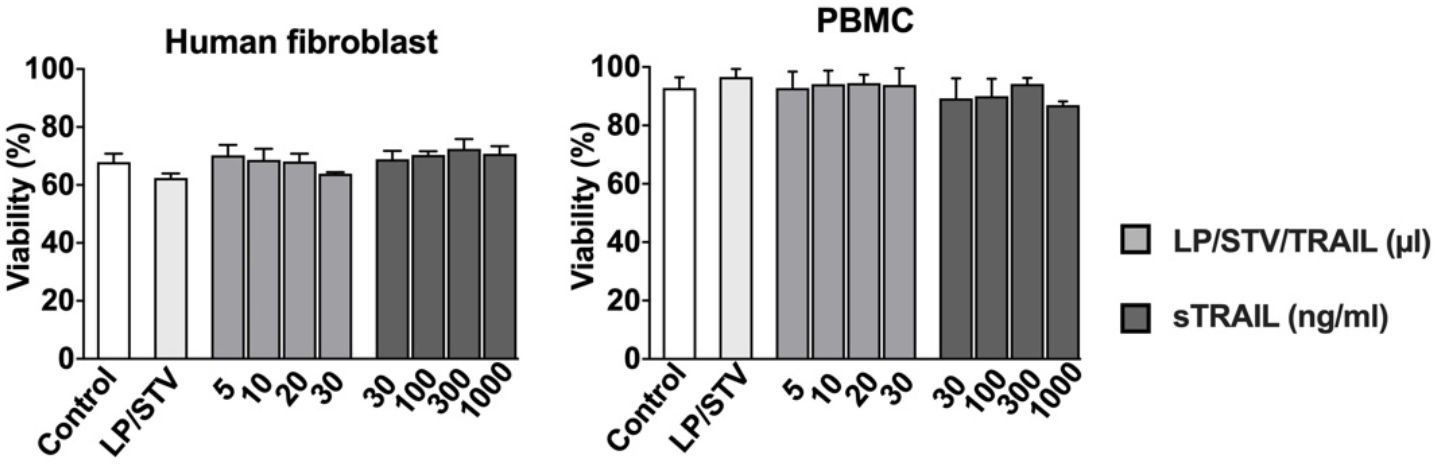
TRAIL-conjugated liposomes are not toxic to non-malignant cells. Non-malignant primary cells (fibroblasts and peripheral blood mononuclear cells (PBMCs)) were incubated with 5-30 μl of TRAIL-conjugated liposomes (LP/STV/TRAIL), 30 μl streptavidin-conjugated liposomes (LP/STV) as negative control and 30-1000 ng/ml soluble recombinant human TRAIL (sTRAIL) for 24 h. Apoptosis was detected using Annexin V-FITC staining. Each bar represents the mean percentage of live cells ± standard deviation of at least three independent repeats. No difference in cell death was observed between control and treated cells (one-way ANOVA followed by Tukey’s test).

NK cells can discriminate between normal and aberrant cells, such as tumor cells, via receptors recognizing stress markers and via their CD16 receptor (FCGR3A) that recognizes antibody-opsonized abnormal cells and trigger antibody-dependent cellular cytotoxicity (ADCC).^1^ To mimic this function of NK cells, we used an engineered immunoglobulin Fc binding peptide (FCP) with high affinity to the Fc region of human antibodies.^12,19^ To verify antibody binding affinity of the FCP, FCP was immobilized in an ELISA plate. Following incubation with a fluorescently-labelled antibody, binding of the antibody to FCP was quantified by spectrophotometry. High fluorescence signal detected in the wells incubated with FITC-Ab in the presence of FCP confirmed the antibody binding capacity of FCP (Supplementary Table 1).

To verify specific, FCP-mediated antibody binding to the liposomes, LP/STV with or without FCP-functionalization were incubated with FITC-Ab, and antibody-binding capacity was monitored by measuring the fluorescence intensity of the liposomes by flow cytometry. FCP functionalization of LP/STV substantially increased their antibody-binding capacity, indicated by their enhanced FITC fluorescence. Additionally, LP/STV functionalized with a higher concentration (30 μg /ml) of FCP showed further increase in FITC fluorescence, (Supplementary Figure 6A), thus 30 μg /ml FCP was chosen as the FCP concentration for the rest of the experiments.

To examine the targeting ability of FCP/antibody-functionalized liposomes, LP/STV/FCP were incubated with an anti-CD123 monoclonal antibody (LP/STV/FCP/CD123), followed by labelling with CM-DiI for traceability. LP/STV/FCP/CD123 showed strong binding to CD123-expressing AML cells, KG1a cells, verifying the targeting potency of liposomes functionalized with specific antibodies (Supplementary Figure 6B).

To generate NK.NPs, liposomes functionalized with TRAIL (LP/STV/TRAIL) and liposomes functionalized with antibodies (LP/STV/FCP/Ab) were fused using a membrane fusion technique^20,21^. To verify liposome fusion, LP/STV/FCP were functionalized with a FITC-tagged antibody. After membrane fusion, TRAIL, on the surface of the fused liposomes was detected with an APC-tagged anti-TRAIL antibody and the proportion of LPs dual positive for FITC and APC were determined with flow cytometry. The results demonstrated that more than 90% of the liposomes became double positive, confirming successful membrane fusion (Figure 1C). NanoSight measurements showed that the fused liposomes (NK.NPs) had an average diameter of 145.2 nm (Supplementary Figure 7) and that the majority of NK.NPs were smaller than 150 nm (D10: 117.3 nm, D50: 145.0 nm and D90: 171.7 nm). DLS measurements and electron microscopy study demonstrated no aggregation occurred during the modification steps (Figure 1) and the PDI was 0.197.

### Natural killer cell-mimic nanoparticles can potently kill AML cells *in vitro*

Acute myeloid leukemia is a very aggressive cancer with a poor clinical outcome. The major challenge in its treatment is drug resistance, immune editing leading to impaired tumour immune surveillance and high interpatient heterogenicity.^23^ With the availability of a safe and well-tested therapeutic antibody (anti-CD38, daratumumab) against AML, we decided to test whether NK.NPs can overcome these challenges.^24^ The efficacy of the NK.NPs to target specific cells via the FCP-conjugated antibody and then kill them, we functionalized the surface of liposomes with the antibody therapeutics, daratumumab, a human anti-CD38 monoclonal antibody used of the treatment of AML, as AML cells often express high levels of CD38 on their surface.^22^ AML cell lines which express CD38 (ML-1, ML-2, Kasumi-1 and OCI-AML2) as well as primary, patient-derived AML samples were tested. Cytotoxicity assays showed that LP/STV/TRAIL had a superior cytotoxic potential than sTRAIL. Furthermore, conjugation of anti-CD38 antibody enhanced the cytotoxic potential of the NK.NPs against one of the CD38-expressing AML cell lines, ML-1 (Figure 4A). Additionally, the cytotoxic potential of the NK cell mimic NPs was also compared to that of activated NK cells, using the NK cell line, KHYG-1. The NK.NPs showed consistently high activity against all AML cell lines tested, while KHYG-1 cells had varying activity and showed poor efficacy against ML-1 (Figure 4A) and ML-2 cells (Figure 4B).

**Figure 4.**
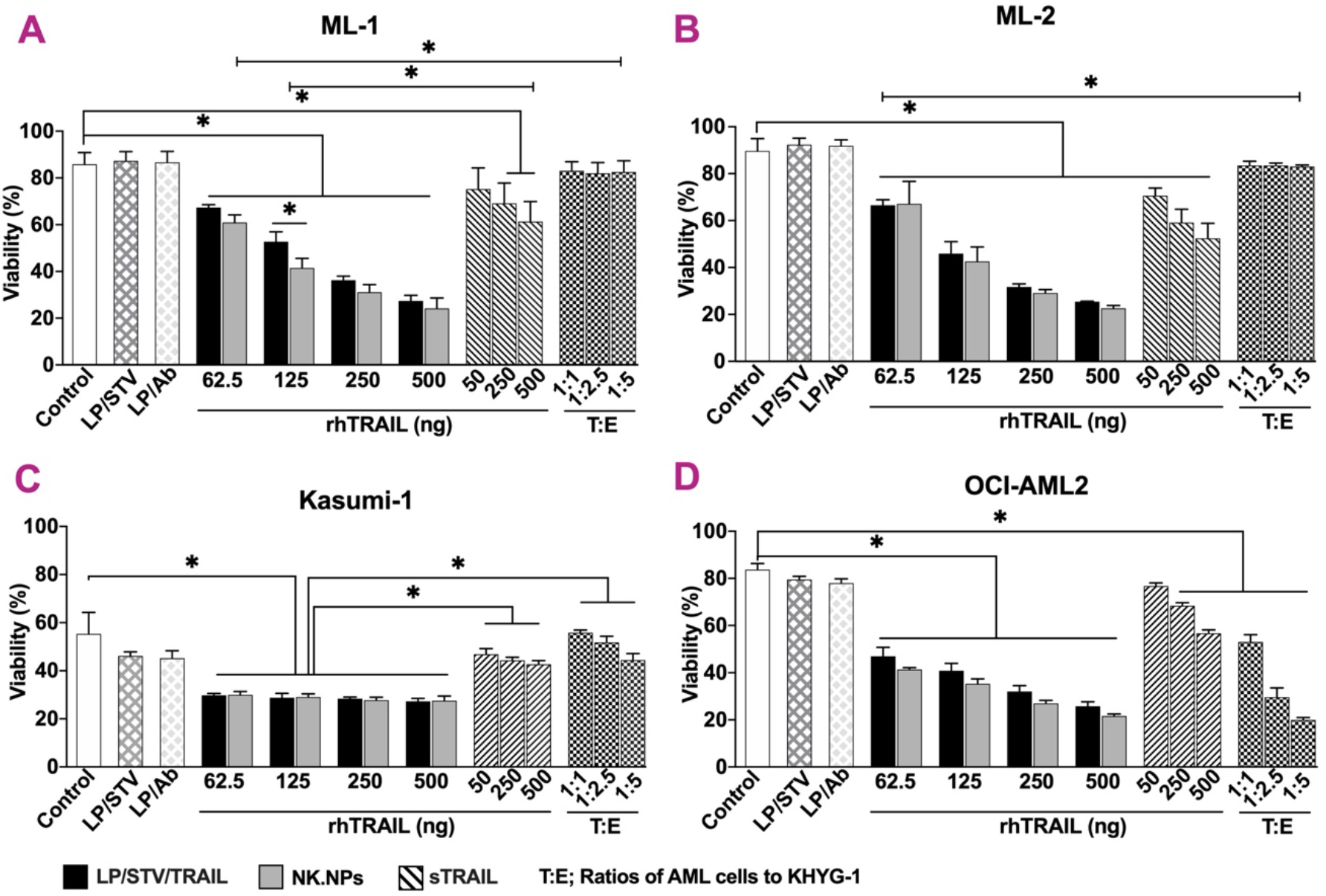
NK cell-mimic nanoparticles are more efficient than NK cells and sTRAIL on myeloma cell lines. Cancer cell lines (as indicated above each graph, ML-1 (A), ML-2 (B), Kasumi-1 (C), and OCI-AML2 (D)) were treated with 62.5-500 ng of TRAIL conjugated liposomes (without antibody, black bars), 62.5-500 ng of TRAIL conjugated liposomes fused to liposome functionalized with anti-CD38 antibody (NK.NPs, grey bars), 20 μl streptavidin-conjugated liposomes (LP/STV, without TRAIL or FCP or antibody as first negative control), 20 μl LP/STV functionalized with antibody (LP/Ab, without TRAIL as second negative control), 50-500 ng soluble recombinant human TRAIL (dashed bars) or were cocultured with CFSE-KHYG-1 cells (effector cell) at 1:1-5 target to effector (T:E) ratios for 24 h. Cell death was detected using Annexin V staining. Each bar represents the mean ± standard deviation of three independent repeats (n=3). The statistical significance was evaluated using one-way ANOVA. P<0.05 was considered significant.

Since primary cancer cells often show higher drug resistance, the efficacy of anti-CD38-functionalised NK.NPs were next tested against primary AML blasts (Figure 5). Functionalization of the NK.NPs with anti-CD38 antibody enabled targeting of CD38-positive AML blasts resulting in enhanced cytotoxicity (Figure 5A and Supplementary Figure 8).

**Figure 5.**
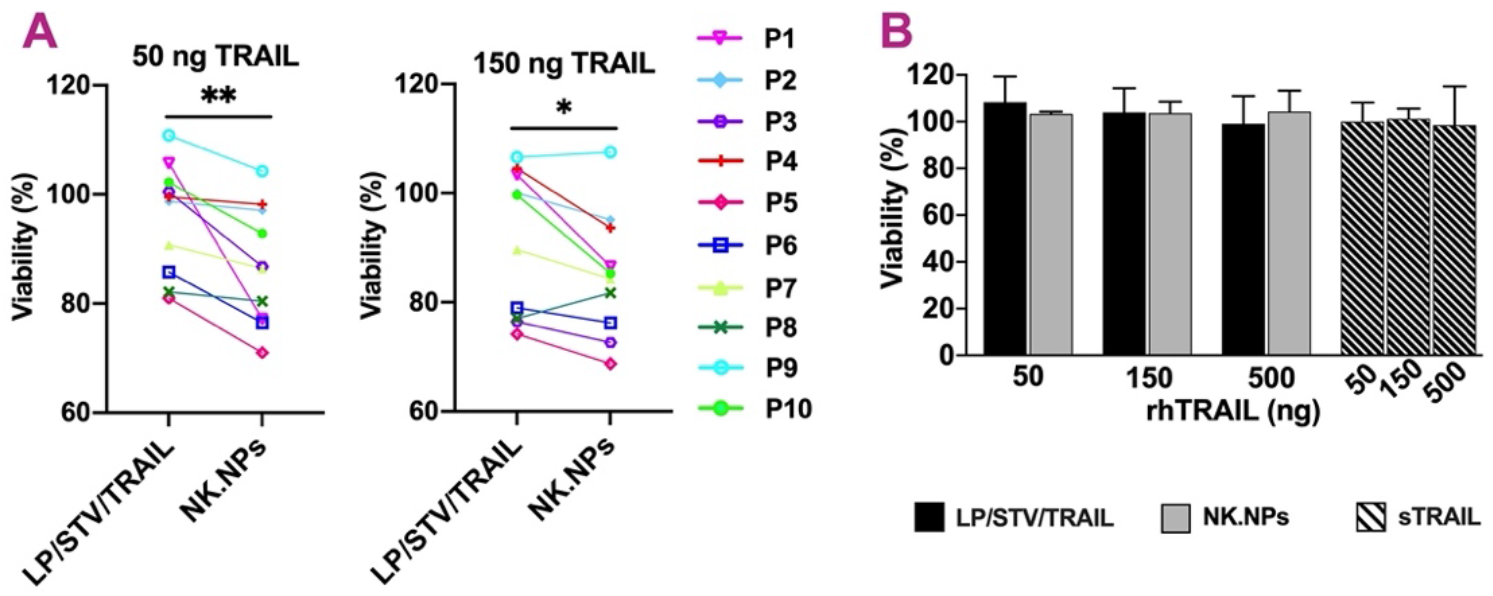
NK cell-mimic nanoparticles could target primary AML blasts *ex vivo*. (A) Primary AML blasts were isolated from 10 patients (P1-10) then were treated with 50 ng or 150 ng of TRAIL-conjugated liposomes (without targeting, LP/STV/TRAIL) or 50 ng or 150 ng of anti-CD38-functionalized NK cell mimic liposomes (CD38-NK.NPs, targeted) for 24 h. Cell death was measured with TO-PRO-3 staining and flow cytometry. (B) Non-malignant primary peripheral blood mononuclear cells from 4 healthy donors were treated with 50-500 ng of LP/STV/TRAIL (grey bars), 50-500 ng of CD38-NK.NPs (black bars), as negative control and 50-500 ng soluble recombinant human TRAIL (sTRAIL) for 24 h. TO-PRO-3 staining was used to identify dead cells. Bars show the percentage of viable cells, normalized to untreated control after 24 h of treatment. Each error bar represents the mean ± SD. * p<0.05, **P < 0.005 determined with paired two-tailed Student’s t-test.

Since drug resistance and relapse in AML are tightly linked and driven by low-differntiation status leukemia-initiating cells, we next assessed the ability of the NK.NPs to eliminate myeloid precursor cells using a colony formation assay. Human primary AML cells were treated with sTRAIL, LP/STV/TRAIL or CD38-NK.NPs for 16 hours after which the AML cells were seeded in Methocult semisolid medium and cultured for 11-13 days to allow growth of progenitor cell-derived AML cell colonies. Unlike sTRAIL, both LP/STV/TRAIL and CD38-NK.NPs blocked colony formation (Figure 6). The effect of CD38-NK.NPs on colony growth inhibition was notably more than LP/STV/TRAIL, corroborating that active targeting enhances efficacy likely by enabling simultaneous activation of higher number of TRAIL receptors.

**Figure 6.**
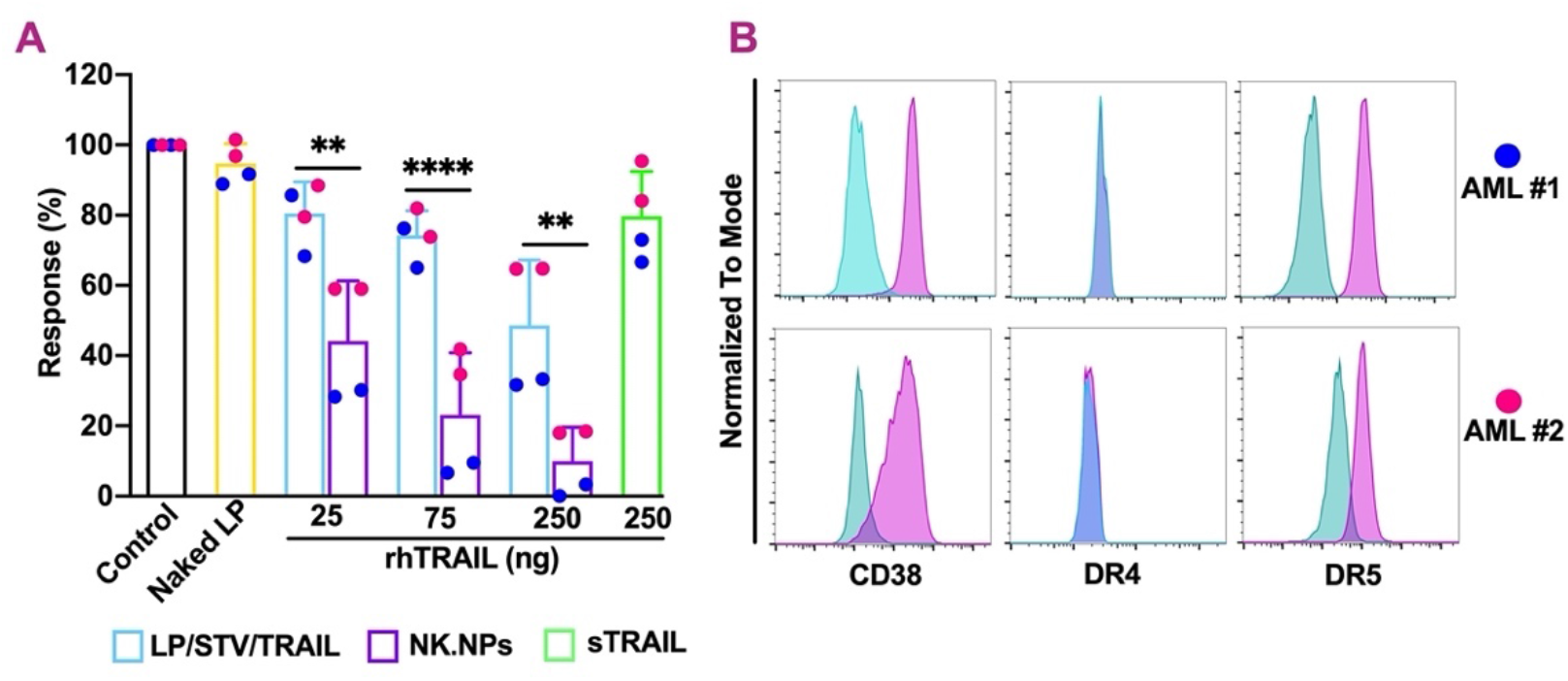
NK cell-mimic NPs inhibit proliferation in primary human AML cells. (A) Primary AML cells isolated from mice bearing patient xenografts were treated with TRAIL-conjugated liposomes (LP/STV/TRAIL, blue bars), anti-CD38 antibody-conjugated NK cell-mimic liposomes (CD38-NK.NPs, purple bars), 10 μl naked liposomes (yellow bar) as negative control and 250 ng soluble recombinant human TRAIL (sTRAIL, green bar) overnight followed by plating in semi-solid medium. Colonies were enumerated after 11-13 days for each xenograft (patient AML#1, blue circles; patient AML#2, red circles) of incubation. (B) Histograms showing CD38-, DR4- and DR5 cell surface expression (pink) and unstained controls (turquoise) for the respective xenografts. Conditions were performed in duplicate. Data are presented as means ± SD. **P < 0.005; ***P < 0.0005; ****P < 0.0005. Significance was determined using one-way ANOVA.

### *In vivo* efficacy of NK cell mimic nanoparticles against acute myeloid leukemia

To assess t*he in vivo* efficacy and potential off-target toxicity of NK.NPs, an AML xenograft model was established by intravenously injecting CD38-positive OCI-AML2 cells into the tail vein of NSG mice. Tumor burden was monitored by measuring the percentage of human CD45^+^ cells in the peripheral blood at various times by collecting blood from the tail vein (Supplementary Figure 9). Circulation of transplanted OCI-AML2 cells became detectable from day 7 and as expected for a disseminated AML, and the majority of human CD45^+^ cells accumulated in the bone marrow.

To assess potential toxic effects of NK.NPs, healthy NSG mice were injected with 2 doses of CD38-NK.NPs or with vehicle (PBS) intraperitoneally, 48 hours apart (Supplementary Figure 10A).). Mice were sacrificed 48 hours after the second injection. The body weight and behavior of the mice were monitored daily. No signs of toxicity, such as appearance, neurologic symptoms, limb weakness, diarrhea or change in body weight were observed. Toxicity to the pancreas, liver, and kidneys was monitored by measuring glucose, liver enzymes, blood urea nitrogen (BUN) and creatinine in the blood, respectively. All these parameters remained within the normal range, confirming that NK.NPs do not cause off-target toxicity (Supplementary Figure 10).

To determine the tumor-targeting potential of NK.NPs *in vivo*, NSG mice with established AML xenografts (4 days post OCI-AML2 injection) were divided into 4 treatment arms and treated with 5 doses of 1) naked liposomes as vehicle, 2) LP/STV/TRAIL, 3) CD38-NK.NPs, or 4) sTRAIL intraperitoneally administered every 48 hours (Figure 7A). Body weight was monitored during the experiment and they were not significantly different among the four treatment groups confirming that NK.NPs had no significant toxic side effects (Figure 7B).

**Figure 7.**
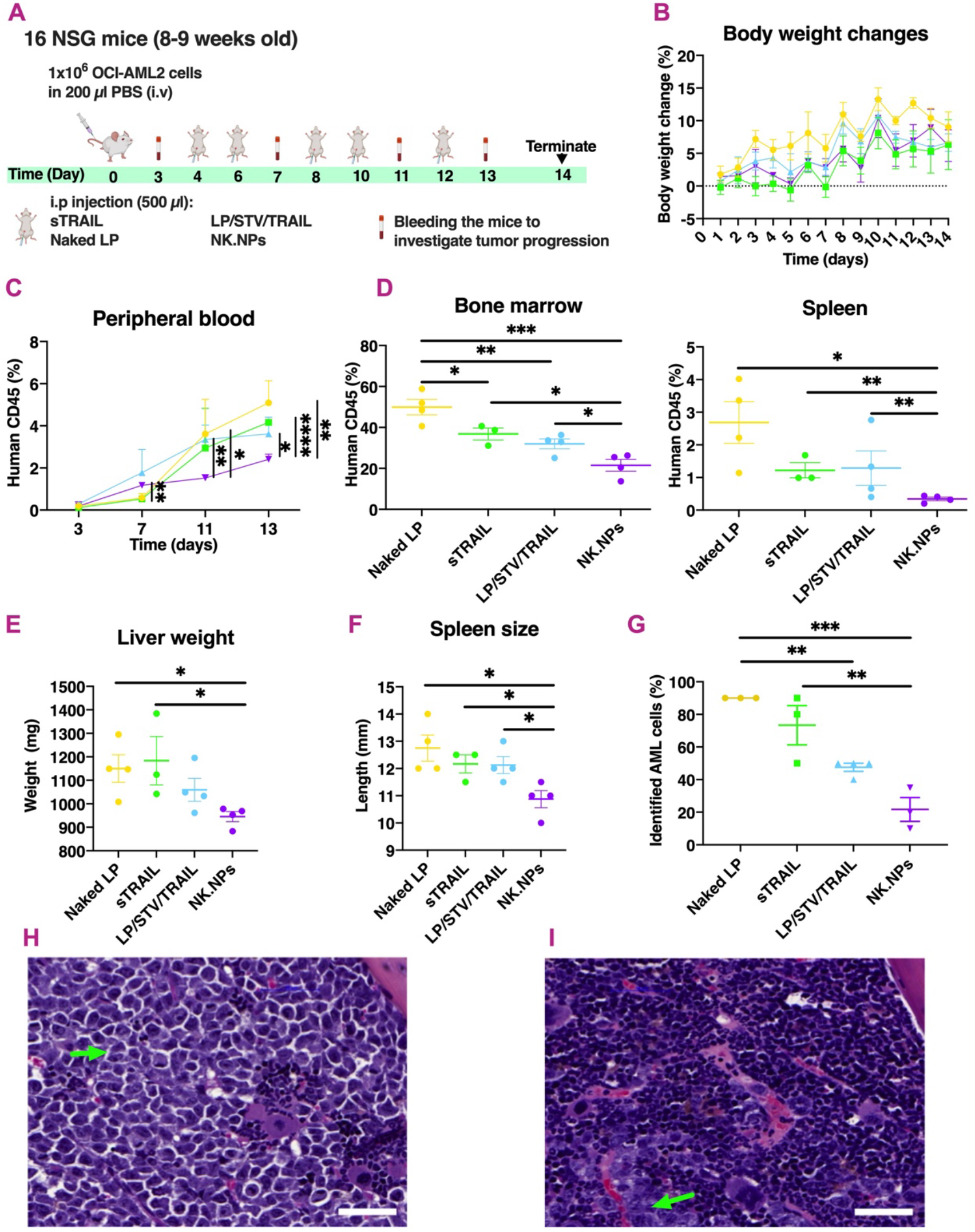
NK cell-mimic nanoparticles can target human AML cells *in vivo*. (A) Schematic illustration of the treatment schedule and sample collection. Healthy, 8-9 weeks old female NSG mice were injected intravenously (i.v) with 1×10^6^ OCI-AML2 cells. Upon establishment of AML, mice were injected intraperitoneally (i.p) with 500 μl of either naked liposomes (Naked LP, indicated in yellow) as vehicle control, soluble recombinant human TRAIL (sTRAIL, green dots), liposomes functionalized with TRAIL (LP/STV/TRAIL, blue dots) or NK cell-mimic NPs (NK.NP, indicated in purple). (B) Changes in body weight. (C and D) Flow cytometric quantification of live human CD45^+^ cells in peripheral blood at different time points during treatment (C). (D) Percentage of live hCD45^+^ cells in the spleen and the bone marrow at the endpoint of the treatment schedule. (E and F) liver weight (E) and spleen size (F). (G) Quantification of AML blast frequencies in hematoxylin and eosin (H&E)-stained slides of tibia from mice at the end of the treatment schedule. (H and I) Representative H&E stained tibia from mouse treated with naked liposome (H) or NK.NP (I). Green arrows point to AML blasts (Magnification= 20X, scale bar = 50 μm). Data are displayed as mean ± SEM. Significance was determined using two-tailed student’s t-test. *P < 0.05; **P < 0.005; ***P < 0.0005; ****P < 0.0005.

Disease progression was monitored by measuring the frequency of OCI-AML2 cells in the peripheral blood (by detecting human CD45^+^ (hCD45^+^) cells) at regular time points. The spread of the disease was determined at the end of the treatment schedule by measuring accumulation of human AML cells (Supplementary Figure 11) in the spleen and bone marrow.

Importantly, treatment with CD38-NK.NP, but not with sTRAIL or LP/STV/TRAIL, reduced the number of CD45^+^ AML cells in the peripheral blood. In the bone marrow and the spleen, where AML cells preferentially accumulate, all treatments were able to reduce the amount of leukemic cells, but the NK.NPs were the most effective, proving their ability to actively target the CD38-expressing AML cells (Figure 7C, 7D). Quantification of hematoxylin and eosin (H&E) stained sections of tibia further confirmed the flow cytometry data, where there was no significant difference between mice injected with naked liposomes and sTRAIL, while NK.NPs and LP/STV/TRAIL dramatically reduced the AML cells accumulation in the bone marrow (Figure 7G-I). Corroborating the ability of CD38-NK.NPs to limit the spread of CD38-positive AML cells, NK.NP-treatment also prevented hepatomegaly and splenomegaly, other symptoms of late stage AML (Figure 7E, 7F).^25^

## Discussion

In this study we developed a nano-formulation which can act as artificial cell-mimics that can replicate key tumor surveillance functions of natural killer (NK) cells: selective tumor recognition and tumor cell killing. To build this NK cell mimic NP, we used PEGylated liposomes as the carrier to avoid clearance by macrophages. In order to replicate the tumor-cell recognition a killing function of NK cells, the surface of the PEGylated liposomes was functionalized with a peptide homologue of the CD16/FCGR3A of NK cells and with the death ligand, TRAIL. For functionalization, we designed a modular assembly of the NK.NP where the functional moieties were conjugated without any chemical modification and the targeting properties were adjustable to suit future personalized therapies.

In the last three decades, considerable efforts have been made to improve the antitumor activity of TRAIL.^8^ Although TRAIL, as a soluble recombinant protein showed good antitumor activity *in vitro* and in preclinical studies, the clinical trials were unsuccessful because of the short half-life time and limited biological activity of sTRAIL.^26^ Subsequently, several studies demonstrated that the cytotoxic activity of TRAIL is enhanced by conjugating it to nanoparticles^5,8^, and a study by Miguel and colleagues have illustrated that TRAIL has to accumulate in the tumor tissue in large amounts to achieve effective tumor eradication, emphasizing that active, targeted delivery of TRAIL to the tumor in its native orientation and conformation is critical to achieve potent activity.^27^ For this reason, we have employed a new functionalization method that allowed immobilization of TRAIL and tumor marker-targeting antibody in an oriented manner without exposing and potentially damaging TRAIL or the antibody by a chemical treatment, such as carbodiimide- or maleimide linkage requiring the reduction of cysteine-SH groups. This is especially important for TRAIL, where cysteine at position 230 is essential for TRAIL’s biological activity.^28^ Although some of the previous studies added an extra cysteine residue to the TRAIL molecule, none could guarantee that during the conjugation reaction the essential Cys^230^ remained intact.^29,30^

In agreement with other studies, TRAIL conjugated to NK.NPs had a robust cytotoxic activity against a broad panel of cancer cell lines of varying sTRAIL sensitivity. In some cases, NP-conjugated TRAIL could also overcome resistance to sTRAIL (e.g. U2OS cells) without being toxic to non-malignant cells. As high sTRAIL concentration is easy to achieve *in vitro* and there are no other somatic cells that may sequester sTRAIL via their decoy receptors (DcR1, DcR2, osteoprotegerin),^31^ active targeting of the NK.NPs in *in vitro* experiments provided limited benefit and only increased the cytotoxicity of the NK.NPs in one cell line. On the other hand, active targeting of TRAIL to the tumor cells became beneficial *ex vivo*, in primary AML blast, showing that when the complexity of the system increases (due to the presence of other, non-malignant mononuclear cells) and TRAIL sensitivity of primary tumor cells being lower, delivery of high TRAIL concentration directly to the surface of tumor cells enhances potency.

Most importantly, in the disseminated AML xenograft model, the NK.NPs not only maintained their activity in an *in vivo* setting, but their advantage over both sTRAIL and liposome-linked TRAIL became apparent. Interestingly, NK.NPs did not only inhibit tumor burden in the bone marrow and spleen more than sTRAIL and LP/STV/TRAIL, but NK.NPs were the only treatment that could reduce the amount of circulating tumor cells in the peripheral blood (Figure 7). This finding also indicates that NK cell-mimics can also target circulating tumor cells and thus might prevent metastasis of solid tumors. Importantly, systematic administration of NK.NPs did not show any detectable organ toxicity, supporting the potential safety of an NK.NP-based therapy. Besides, NK cell-mediated killing has been shown to be able to induce and activate adaptive immune cells through stimulating antigen-specific CD4^+^ T-cell responses, thus promoting antigen cross-presentation to CD8^+^ cytotoxic T.^35^ Therefore, NK.NPs-mediated killing might synergize with other immune cells.

Recently, in another study^32^ liposomes were functionalized with TRAIL in combination with E-selectin, thereby enabling the liposomes to latch onto leukocytes and thus get delivered to the tumor cells. These liposomes (functionalized with 50 μg TRAIL, which is similar to the concentration in this study) could also efficiently eliminate circulating tumor cells. This formulation however had a limited ability to target tumor cells at the primary tumor site,^32^ which may indicate that immune cell-based, indirect NP targeting may be hampered when immune cells are inactivated in cancer patients due to tumor immune editing or active immunosuppression of immune effectors. Consequently, NP formulation that can overcome this limitation are predictably more potent.^33,34^

In summary, the developed NK cell-mimic nanoplatform is unique because this design enables conjugation of the functional moieties (TRAIL and tumor-targeting antibody) to the NPs in the correct orientation and intact conformation. Furthermore, by using FCP functionalization of the NPs, a modular system was created where the NK.NPs can be decorated with any desirable therapeutic antibody matching the biomarker profile of individual tumors and thus applicable for personalized therapy.

### Conclusions

This study developed an NK cell-mimic nanoparticle (NK.NP) to target and recognize tumor cells, followed by tumor cell killing without any obvious systemic toxicity in an *in vivo* mouse model of AML. Importantly, the targeting strategy used in the design of the NK.NPs improved the biological activity of TRAIL, thus enabling efficient tumor cell elimination and reduction of AML development *in vivo*. A second important feature of the NK.NPs is its modular built, where the NP can be functionalized by any tumor-targeting antibody via the FCP immunoglobulin-binding peptide.

Although the described NK.NPs only mimic the surface characteristics of NK cells, they can be further improved in the future by encapsulation of NK cell-produced cytolytic proteins, such as perforin and granzymes, to also mimic the cell lysis function of NK cells. In addition, the NK.NPs can be loaded with small molecule drugs to overcome TRAIL sensitivity, thus providing a highly efficient therapeutics suitable for a tailor-made, personalized therapy.

## Supporting information

Supplementary Figure 11. Gating strategy used for assessing AML tumour burden in vivo.

Supplementary Table 1. Immunoglobulin binding peptide have binding affinity to human antibody.

Supplementary Figure 10. NK cell-mimic NPs do not have significant toxic side effects.

Supplementary Figure 9. Human AML cell lines successfully engrafted in the NSG mice.

Supplementary Figure 8. Correlation between CD38 expression and targeting potential of NK cell-mimic nanoparticles on primary AML blasts in vitro.

Supplementary Figure 7. Characterization of NK cell-mimic nanoparticles with NanoSight.

Supplementary Figure 6. Functionalization of liposomes with immunoglobulin binding peptide and antibody.

Supplementary Figure 5. Flow cytometric analysis of liposome phagocytosis by macrophages.

Supplementary Figure 4. Optimization of TRAIL conjugation to maximize TRAIL functionalization of liposomes.

Supplementary Figure 3. Functionalization of liposomes with Strep-TRAIL.

Supplementary Figure 2. Liposome surface modification with streptavidin.

Supplementary Figure 1. Purification of Strep-TRAIL expressed in E. coli BL21 bacteria.

## Acknowledgements

We would like to acknowledge the scientific and technical assistance of the University of Galway Flow Cytometry and the Centre for Microscopy and Imaging core facilities, and the University of Bern Flow Cytometry and Animal facilities. We thank the team of the Translational Research Unit (TRU) of the Institute of Pathology for their technical support. We also thank Coline Nydegger, Kristyna Hlavackova, Oceane Derivaz, and Vivian Pham Vu for their kind helps with *in vivo* studies at the University of Bern. We are grateful to Dr. Éadaoin Timmins (University of Galway) for her help with the TEM study, to Prof. Christoph von Ballmoos and his student Lukas Rimle for providing lab facilities to generate NK.NPs for the *in vivo* studies at the Department for Chemistry, Biochemistry and Pharmacy of the University of Bern. We thank Ursina Luethi (University of Bern) for advice and technical help with the colony forming assay. We also thank Dr. Michael Rainy (University of Galway) for his technical help with protein purification.

