## Supplementary Figure 11. Gating strategy used for assessing AML tumour burden in vivo. for "Natural killer cell-mimic nanoparticles can actively target and kill acute myeloid leukemia cells"

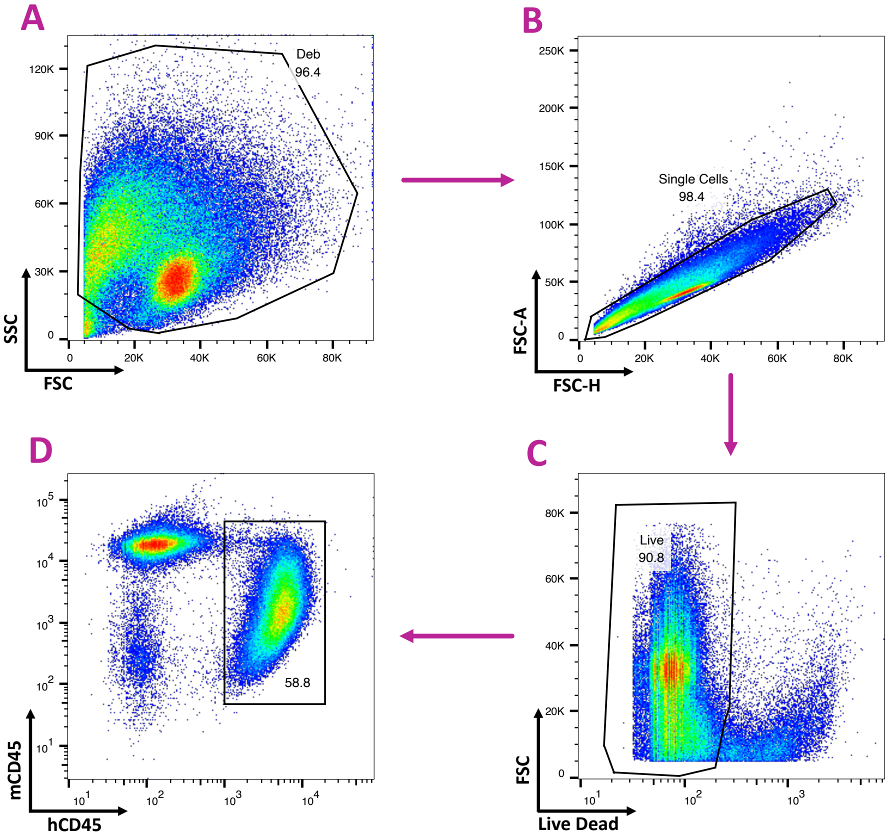


**Supplementary Figure 11. Gating strategy used for assessing AML tumour burden *in vivo*.** First, in a side versus forward scatter plot (FSC vs SSC), cell debris was excluded (A), followed by gating on single cells by plotting forward scatter area versus forward scatter height (B, FSC-A vs FSC-H). Afterwards, living cells were gated by eliminating dead cells stained with live/dead fixable blue dead cell stain (C). Finally, live cells plotting using anti-mouse CD45 (mCD45) versus anti-human (hCD45) fluorochrome-conjugated antibodies allowed identification of donor AML cells (hCD45$^{+}$) (D).
