## Supplementary Table 1. Immunoglobulin binding peptide have binding affinity to human antibody. for "Natural killer cell-mimic nanoparticles can actively target and kill acute myeloid leukemia cells"

| Ratio | Flu. Int. Mean (n=3) | Flu. Int. Change | Flu. Int. change VS undiluted |
|---|---|---|---|
| **0** | 1500176 | 0 | - |
| **1 (6 μg)** | 1852520 | 352344 | 1 |
| **1/3** | 1847751 | 347575 | 0.9865 |
| **1/10** | 1838792 | 338616 | 0.9610 |
| **1/30** | 1823269 | 323093 | 0.9170 |
| **1/100** | 1672276 | 172100 | 0.4884 |
| **1/300** | 1594968 | 94792 | 0.2690 |
| **1/1000** | 1530827 | 30651 | 0.0870 |
| **1/3,000** | 1515462 | 15286 | 0.0434 |
| **1/10,000** | 1510440 | 10264 | 0.0291 |
| **1/30,000** | 1512639 | 12463 | 0.0354 |

Fluorescence intensity changes (Flu. Int. Change) versus the highest concentration (undiluted FCP).
