## Supplementary Figure 10. NK cell-mimic NPs do not have significant toxic side effects. for "Natural killer cell-mimic nanoparticles can actively target and kill acute myeloid leukemia cells"

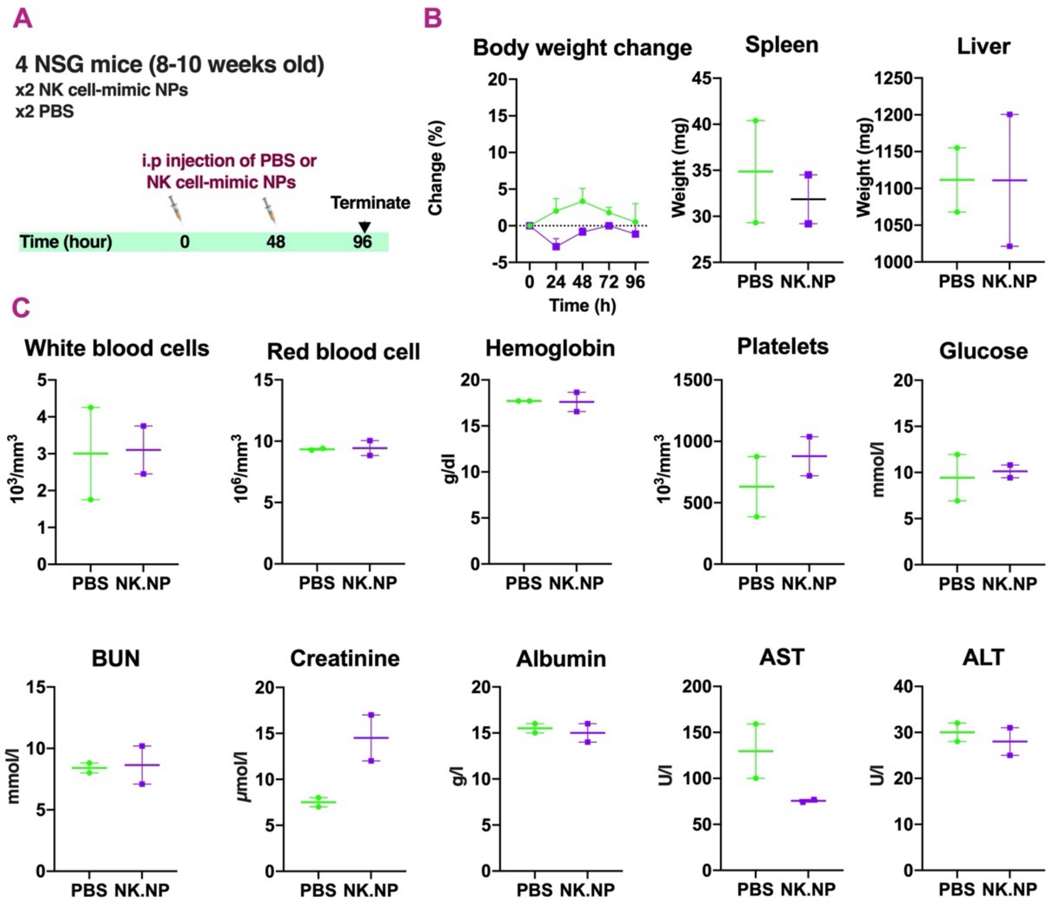


**Supplementary Figure 10. NK cell-mimic NPs do not have significant toxic side effects.** (A) Schematic illustration of the treatment. 8-10 weeks old female NSG mice were injected with 500 µl NK cell-mimic NPs (NK.NP; purple) or with 500 µl PBS (vehicle; green). (B) Changes in body weight, spleen- and liver weight over time. (C) Blood cell count changes and plasma biochemical analysis after 4 days. Cell types and examined plasma markers are indicated at the top of each graph. Data are displayed as mean ± SEM. Significance was determined using a two-tailed Student's t-test. *P < 0.05.
