## Supplementary Figure 9. Human AML cell lines successfully engrafted in the NSG mice. for "Natural killer cell-mimic nanoparticles can actively target and kill acute myeloid leukemia cells"

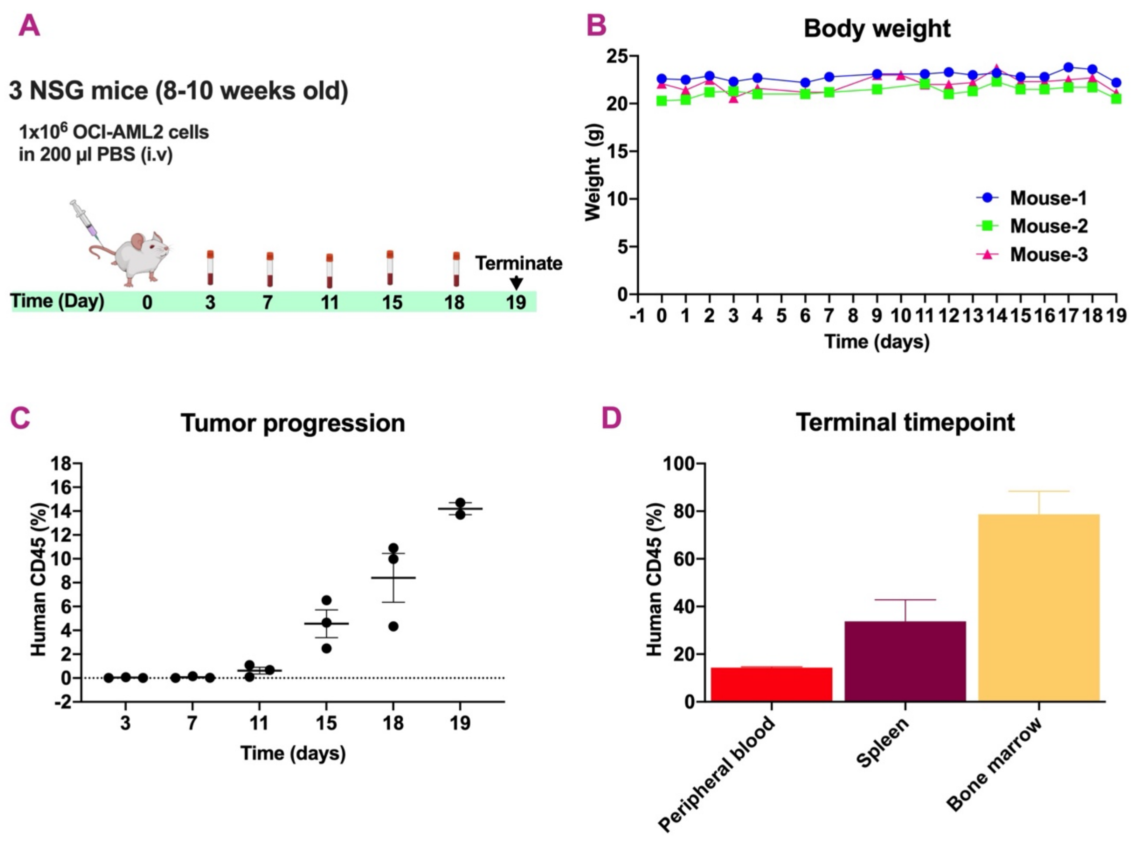


**Supplementary Figure 9. Human AML cell lines successfully engrafted in the NSG mice.** (A) Schematic illustration of the experimental plan. Three healthy 8-10 weeks old female NSG mice were injected intravenously (i.v) with $1\times10^6$ OCI-AML2 cells. (B) Changes in body weight. (C-D) Flow cytometric quantification of live human $CD45^+$ AML cells in peripheral blood at different time points (C), also spleen, and bone marrow at the endpoint of the experiment (D). Each error bar represents the mean ± SD.
