## Supplementary Figure 8. Correlation between CD38 expression and targeting potential of NK cell-mimic nanoparticles on primary AML blasts in vitro. for "Natural killer cell-mimic nanoparticles can actively target and kill acute myeloid leukemia cells"

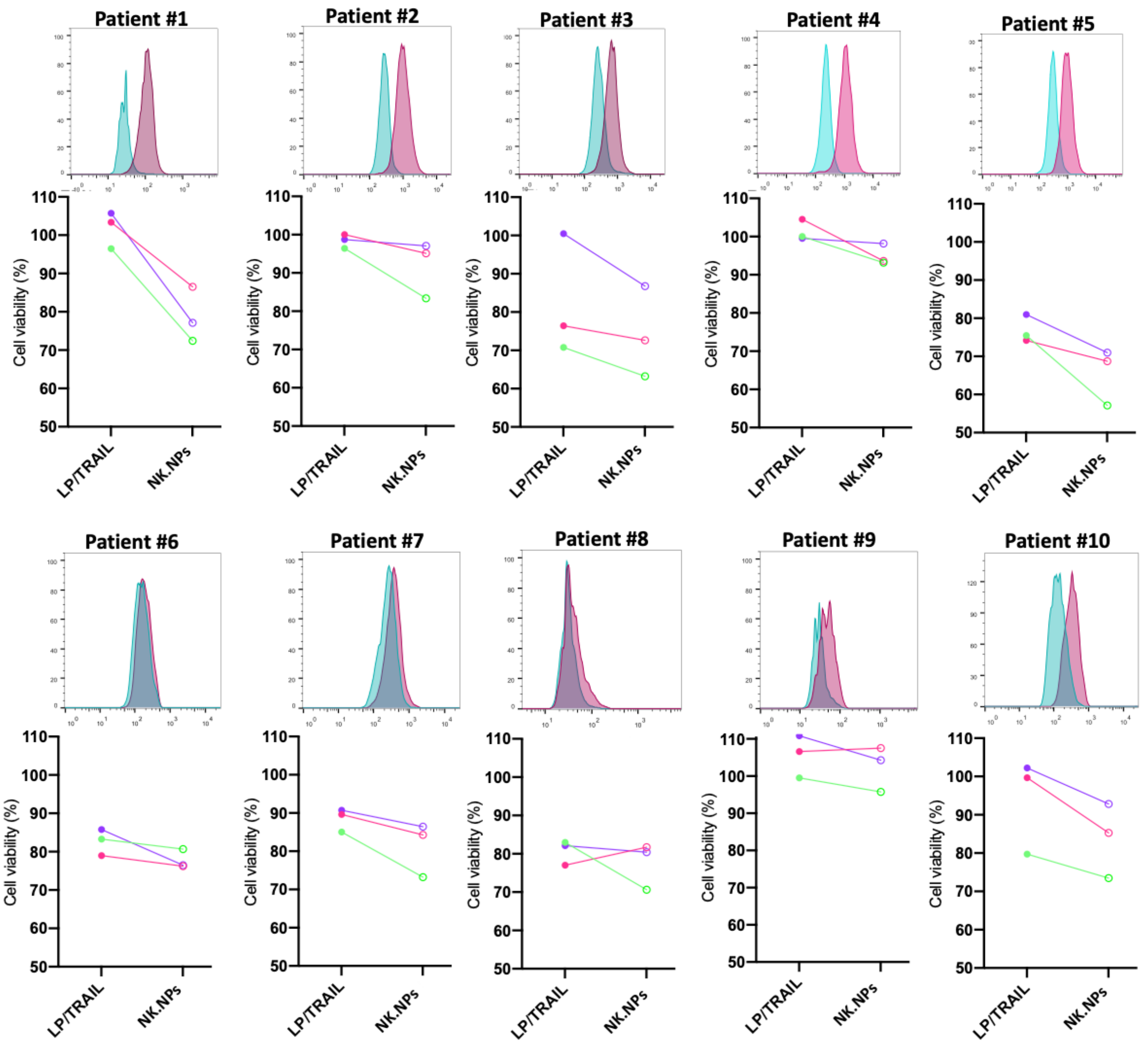


**Supplementary Figure 8. Correlation between CD38 expression and targeting potential of NK cell-mimic nanoparticles on primary AML blasts *in vitro*.** Histograms indicated blast cells isolated from acute myeloid leukemia (AML) patients stained with anti-CD38 antibody (red) and unstained controls (green). Graphs belove each histogram represent AML cells isolated from the same patient which treated with 50 ng (purple lines), 150 ng (pink lines) or 500 ng (green lines) of LP/STV/TRAIL or anti-CD38-conjgated NK cell mimic NPs (NK.NPs). The upper row are AML blasts with high CD38 expression and the lower row with low or no CD38 expression.
