## Supplementary Figure 7. Characterization of NK cell-mimic nanoparticles with NanoSight. for "Natural killer cell-mimic nanoparticles can actively target and kill acute myeloid leukemia cells"

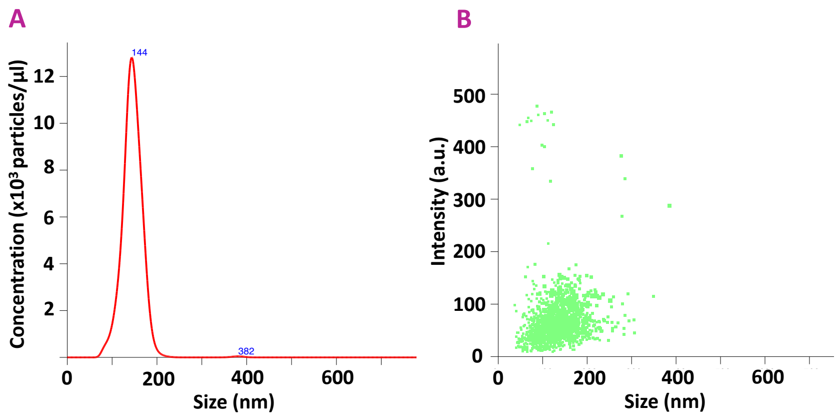


**Supplementary Figure 7. Characterization of NK cell-mimic nanoparticles with NanoSight.** NK.NPs were prepared, and 10 times diluted in PBS. Followed by injecting in a green laser chamber using a NanoSight (Nanosight NS300 system, Malvern). The camera was set in level 12 and capture duration of 60 seconds to capture video with minimal background noise and sufficient contrast to identify particles clearly. The recorded videos were automatically analyzed using nanoparticle tracking analysis (NTA) software, and the size distribution with an estimate of the number of particles were calculated. (A) Histogram represents the size distribution and abundance of NK.NPs that describes the relationship between particle number and size distribution. (B) Scatter plot graph of NK.NPs.
