## Supplementary Figure 6. Functionalization of liposomes with immunoglobulin binding peptide and antibody. for "Natural killer cell-mimic nanoparticles can actively target and kill acute myeloid leukemia cells"

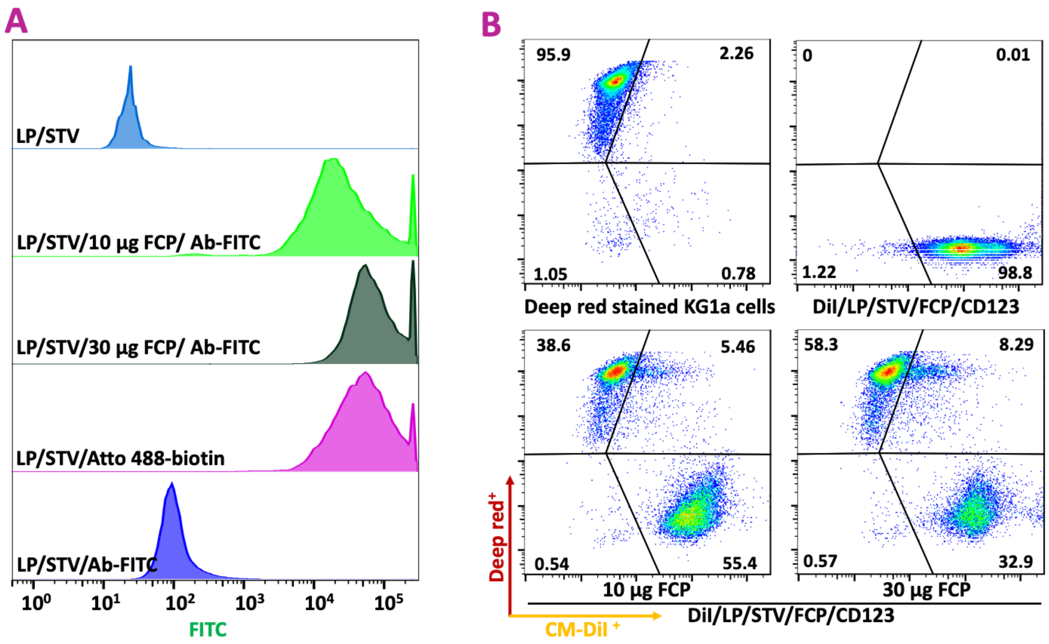


**Supplementary Figure 6. Functionalization of liposomes with immunoglobulin binding peptide and antibody.** (A) Histograms demonstrate the mean fluorescence of liposomes measured with flow cytometry. Liposomes modified with streptavidin (LP/STV) as baseline control, LP/STV incubated with Atto 488-biotin as positive control or with the indicated amount of immunoglobulin binding peptide followed by further incubation with FITC conjugated antibody (LP/STV/10 or 30 µg FCP/Ab-FITC), LP/STV incubated with Ab-FITC as negative control (LP/STV/Ab-FITC). (B) Flow cytometry plots of antibody conjugated liposomes and KG1a cells. liposomes functionalized with 10 µg/ml or 30 µg/ml of FCP were incubated with anti-CD123 antibody and then labelled with CM-DiI (DiI/LP/FCP/CD123 Ab). Next, the pre-seeded deep red stained KG1a cells were incubated with DiI/LP/FCP/CD123 Ab for 90 minutes followed by measuring fluorescence intensities using flow cytometry.
