## Supplementary Figure 5. Flow cytometric analysis of liposome phagocytosis by macrophages. for "Natural killer cell-mimic nanoparticles can actively target and kill acute myeloid leukemia cells"

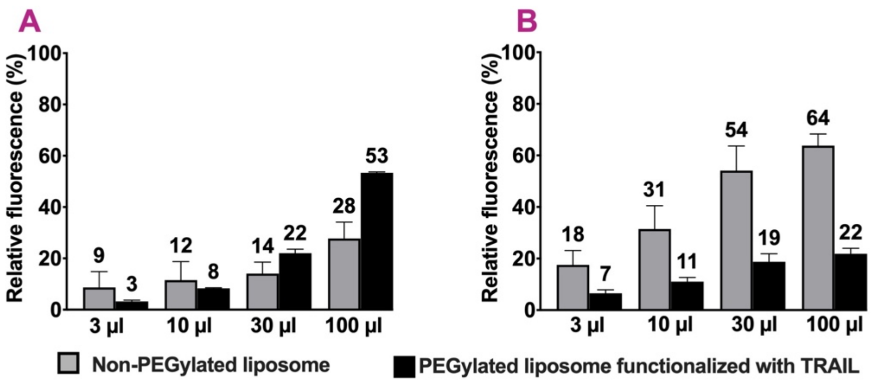


**Supplementary Figure 5. Flow cytometric analysis of liposome phagocytosis by macrophages.** THP-1 derived macrophages were incubated with the indicated volumes and formulations of CM-DiI-labelled liposomes for 3 h; cells stained with APC-tagged anti-CD45 antibody and subjected to flow cytometry measurement. Numbers above bars indicate the average percentages of (n=3) liposomes that escaped from phagocytosis (A) and THP-1 derived macrophages that phagocyted liposomes (B).
