## Supplementary Figure 4. Optimization of TRAIL conjugation to maximize TRAIL functionalization of liposomes. for "Natural killer cell-mimic nanoparticles can actively target and kill acute myeloid leukemia cells"

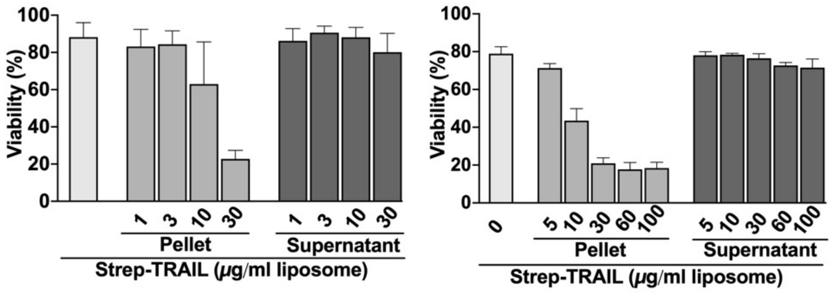


**Supplementary Figure 4. Optimization of TRAIL conjugation to maximize TRAIL functionalization of liposomes.** Colo205 cells were treated with either LP/STV/TRAIL generated by functionalizing LP/STVs with an increasing concentration of Strep-TRAIL or the supernatants of the Strep-TRAIL conjugation reaction for 3 hours followed by quantification of cell death induced using Annexin V-FITC staining.
