## Supplementary Figure 3. Functionalization of liposomes with Strep-TRAIL. for "Natural killer cell-mimic nanoparticles can actively target and kill acute myeloid leukemia cells"

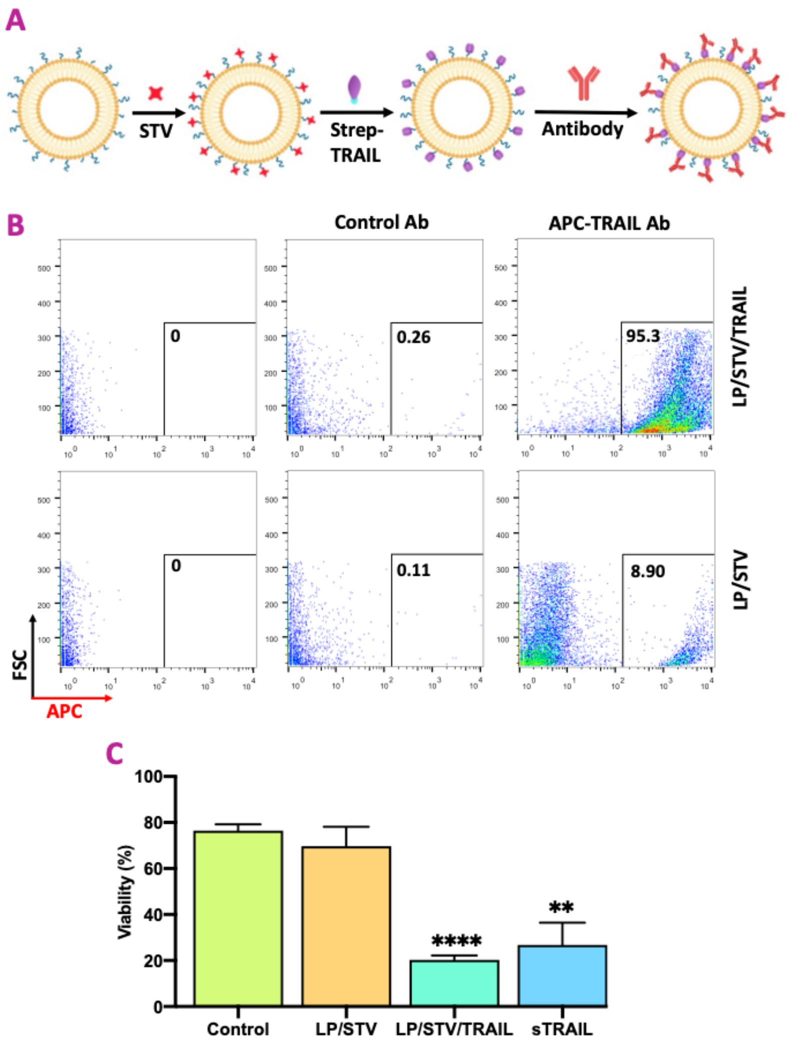


**Supplementary Figure 3. Functionalization of liposomes with Strep-TRAIL.** A) Schematic overview of the surface modification of liposomes with Strep-TRAIL and detection of liposome-bound TRAIL using anti-TRAIL antibody. (B) Detection of TRAIL on liposomes functionalized with streptavidin with flow cytometry (LP/STV: liposome conjugated with streptavidin, LP/STV/TRAIL: LP/STV after conjugation of TRAIL). Liposomes were incubated with either anti-TRAIL antibody (APC tagged) or a nonspecific APC-conjugated antibody (Control Ab) and fluorescence intensity of samples was measured with flow cytometry. (C) Biological/cytotoxic activity of LP/STV/TRAIL. TRAIL-sensitive Colo205 colon carcinoma cells were treated with LP/STV, LP/STV/TRAIL and soluble recombinant human TRAIL (sTRAIL) as a positive control for 3 h after which the number of dying cells was quantified using Annexin V-FITC staining. Each bar represents the mean ± SD of three independent experiments. The statistical significance was determined using a two-tailed t-test. Asterisks denote significant differences between treatments and control; $^{**}P < 0.005$; $^{****}P < 0.00005$.
