## Supplementary Figure 2. Liposome surface modification with streptavidin. for "Natural killer cell-mimic nanoparticles can actively target and kill acute myeloid leukemia cells"

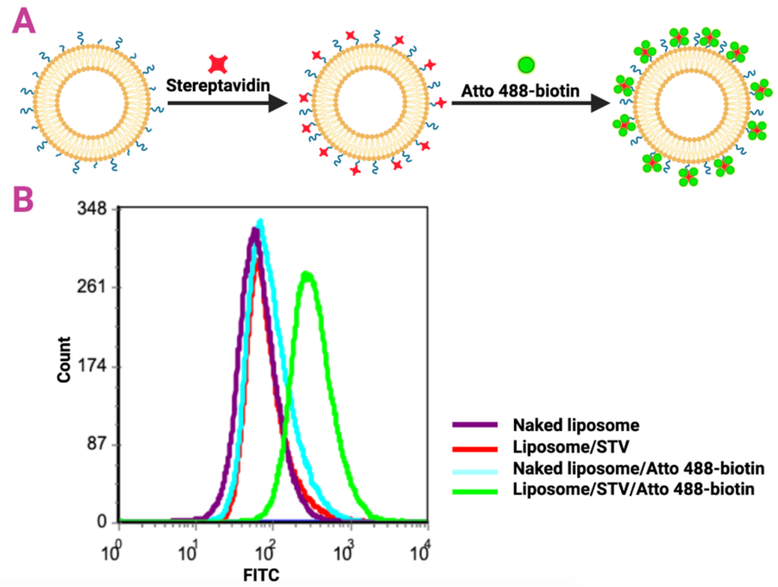


**Supplementary Figure 2. Liposome surface modification with streptavidin.** (A) Schematic overview of the surface modification process of the liposomes with streptavidin (Liposome/STV) and its detection with Atto 488-biotin. (B) Histogram showing the mean fluorescence of liposome formulations (as indicated in the legend) measured with flow cytometry.
