## Supplementary Figure 1. Purification of Strep-TRAIL expressed in E. coli BL21 bacteria. for "Natural killer cell-mimic nanoparticles can actively target and kill acute myeloid leukemia cells"

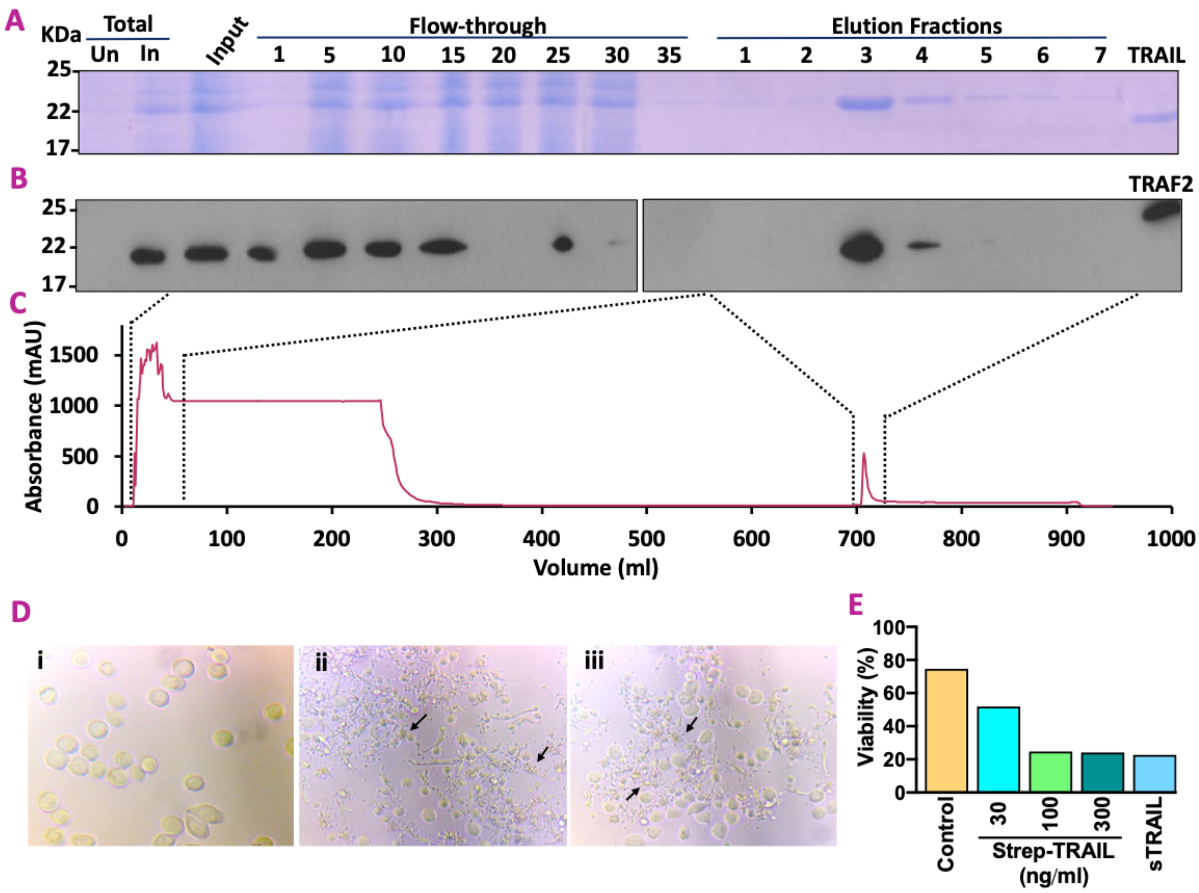


**Supplementary Figure 1. Purification of Strep-TRAIL expressed in *E. coli* BL21 bacteria.** (A-C) Purification of recombinant TRAIL with an N-terminal Strep-tag II. SDS-PAGE analysis using Coomassie blue staining (A) and western immunoblotting (B) at the indicated stages of purification. Total Un: total cell lysate from bacterial culture prior to induction of TRAIL expression (uninduced). Total In: total cell lysate 5 hours after induction of TRAIL expression with IPTG. As a positive control for TRAIL on the Coomassie blue gel, untagged recombinant human TRAIL was used (runs approximately 3 kDa smaller than Strep-TRAIL), and as a positive control for Strep-tag detection with western blotting, Strep-tagged recombinant TRAF2 protein was used (runs at approximately 26 kDa). (C) Representative UV (280 nm) profile of chromatography purification. The cleared bacterial lysate was loaded onto a StrepTrap™ HP affinity column using an AKTA purifier FPLC system, and the chromatogram was recorded as UV absorbance at 280 nm. (D) Morphological assessment of the cytotoxic activity of Strep-TRAIL on Colo205 cells. Light microscopic image of Colo205 cells (i) without treatment, (ii) after treatment with 80 ng/ml of sTRAIL or with (ii) untagged recombinant human TRAIL (sTRAIL) for 3 h. Arrows point to cells with apoptotic morphology. (E) Quantitation of the cytotoxic activity of Strep-TRAIL. Colo205 colon cancer cells were treated with increasing concentration of Strep-TRAIL or sTRAIL as a positive control for 3 h after which the number of dying cells was measured using Annexin V-FITC staining.
